# Zinc-Induced Fluorescence Turn-on in Native and Mutant Phycoerythrobilin-Binding Orange Fluorescent Proteins

**DOI:** 10.1101/2023.08.11.552977

**Authors:** Gary C. Jensen, Makena K. Janis, Jazzmin Jara, Nasir Abbasi, Melissa L. Zastrow

## Abstract

Cyanobacteriochrome (CBCR)-derived fluorescent proteins are a class of reporters that can bind bilin cofactors and fluoresce across the ultraviolet to near-infrared spectrum. Derived from phytochrome-related photoreceptor proteins in cyanobacteria, many of these proteins use a single small GAF domain to autocatalytically bind a bilin and fluoresce. The second GAF domain of All1280 from *Nostoc* sp. PCC7120 is a DXCF motif-containing protein that exhibits blue light-responsive photochemistry when bound to its native cofactor, phycocyanobilin. GAF2 can also bind non-photoswitching phycoerythrobilin (PEB), resulting in a highly fluorescent protein. Given the small size, high quantum yield, and that, unlike green fluorescent proteins, bilin-binding proteins can be used in anaerobic organisms, the orange fluorescent GAF2-PEB protein is a promising platform for designing new genetically encoded metal ion sensors. Here we show that GAF2-PEB undergoes a ∼5-fold reversible zinc-induced fluorescence enhancement with blue-shifted emission maximum (572 to 517 nm), which is not observed for a related PEB-bound GAF from *Synechocystis* sp. PCC6803 (Slr1393g3). Zn^2+^ significantly enhances GAF2-PEB fluorescence across a biologically relevant pH range from 6.0–9.0 and with pH-dependent µM to nM dissociation constants. Site-directed mutants aiming to sterically decrease and increase access to PEB show a decreased and similar amount of zinc-induced fluorescence enhancement, respectively. Mutation of the cysteine residue within the DXCF motif to alanine abolishes zinc-induced fluorescence enhancement. Collectively, these results support the presence of a fluorescence enhancing Zn^2+^ binding site in GAF2-PEB likely involving coordination to the bilin cofactor and requiring a nearby cysteine residue.

## Introduction

Genetically encoded protein-based sensors are invaluable tools for studying biological processes involving various analytes, including metal ions. Many of these biosensors, however, rely on green fluorescent proteins (GFPs) and their derivatives, which are limited to aerobic systems since molecular oxygen is required to form the intrinsic fluorescent chromophore.^1,2^ Recent efforts have led to the engineering of various alternative fluorescent proteins that do not require oxygen to become fluorescent in living systems and which can be applied to anaerobic organisms, like soil or gut bacteria.^3,4^ These proteins often use cofactors such as flavin mononucleotide or heme-derived linear tetrapyrroles (bilins), which bind noncovalently or covalently to form a fluorescent protein.^5–8^ Among these reporters, the flavin-binding fluorescent proteins and near-infrared (NIR) fluorescent proteins are relatively well studied and cover the green and NIR emission ranges of the light spectrum.^5–7^ Engineered monomeric NIR fluorescent proteins (miRFPs) have been developed from bacterial phytochrome photoreceptors and cyanobacteriochromes (CBCRs) that can bind the biliverdin IXα (BV) cofactor.^9–11^ Although the BV chromophore is produced in live cells via heme oxygenase-catalyzed breakdown of heme, the production of fluorescence does not strictly require oxygen because bilin-binding fluorescent proteins can bind exogenously added bilin molecules.^12^ While these bacterial phytochrome-based NIR fluorescent proteins are comprised of two domains, a PAS (Per/Arnt/Sim) and a GAF (cGMP-specific phosphodiesterases, adenylyl cyclases, and FhlA) domain (∼35 kDa combined), CBCRs can use a single ∼18 kDa GAF domain to bind the bilin cofactor.^13,14^ Compared to native bacteriophytochromes, CBCRs from various species of cyanobacteria can bind a variety of linear tetrapyrroles, or bilins, and emit fluorescence at wavelengths from the ultraviolet across the visible spectrum to the NIR.^15^

In contrast to the GAF domains isolated from related canonical phytochromes, independent CBCR GAF domains bound to a bilin chromophore can undergo photoconversion.^13,14^ Bilins typically bind to GAF domains via a C3^1^ linkage between the bilin A ring and a conserved cysteine residue (Scheme S1), and in many CBCRs this binding occurs autocatalytically.^14^ Several bilin cofactors have been found in CBCR GAF domains, including phycocyanobilin (PCB), phycoviobilin (PVB), and phycochromobilin (PΦB).^16–19^ CBCRs use a variety of different color-tuning mechanisms that give rise to their interesting photophysical properties. For example, the third GAF domain of Slr1393 from *Synechocystis* sp. PCC 6803 (UniProtKB P73184, residues 441–596) binds PCB and exhibits photoswitching between green (P_g_: λ_max_ = 648 nm, Φ_F_ = 0.07) and red (P_r_: λ_max_ = 536 nm, Φ_F_ = 0.04) states, as a result of photoisomerization between the 15Z and 15E configurations about the D-ring (Scheme S1).^20–22^ The second GAF domain of All1280 from *Nostoc* sp. PCC7120 (UniProtKB Q8YXD3, residues 562-727) harbors a DXCF motif conserved in some CBCRs that provides an additional cysteine residue near the bilin cofactor.^23,24^ In GAF2, this residue reversibly binds to the C10 position of PCB (some of which is autoisomerized upon binding to PVB), breaking the conjugation of the cofactor and allowing for blue/green photoswitching (Scheme S1). Photoswitching CBCRs have limited utility as fluorescent biomarkers, however, given that they have relatively low fluorescence quantum yields. To overcome this obstacle, the non-photoswitching phycoerythrobilin (PEB) cofactor can be used. PEB is identical to PCB except it lacks the Δ15,16 double bond between pyrrole ring C and D, and when bound to a variety of CBCRs results in fluorescent proteins with high quantum yields and molar extinction coefficients.^25,26^ These CBCR GAFs can be heterologously coexpressed in *E. coli* with the biosynthesis enzymes for PEB production from a heme precursor (heme oxygenase and PebS).^27–30^ Among the CBCR GAFs tested for PEB binding, All1280 GAF2 is one of the brightest PEB binding CBCRs (Φ_F_ = 0.68, ε_495/546_ = 49,000/24,000 M^-1^ cm^-1^) and shows absorption maxima that could make it suitable as an acceptor in Förster Resonance Energy Transfer (FRET)-based platforms utilizing proteins that emit green fluorescence as the donor.^25^

Transition metals like zinc are essential to living organisms due to their roles in catalysis, structural stabilization of proteins, and biological signaling.^31^ Fluorescent sensors are frequently used to study the roles of metals in biology given their ability to visualize real-time metal ion binding in living samples.^32^ Fluorescent protein-based sensors can be genetically encoded into living cells or organisms, allowing for species-specific, cell-specific, or subcellular imaging of metal ions.^33–37^ Fluorescent proteins can be used as sensors by attaching or embedding a metal binding site in a single fluorescent protein that modulates fluorescence in response to the metal ion. Multi-fluorescent protein-based probes can also be prepared by employing a FRET approach where two proteins are linked by a metal site that undergoes a conformational change upon metal binding. Given that most genetically encoded metal sensors are based on GFPs and related mFruits,^33–36^ here we sought to evaluate the PEB-binding orange fluorescent GAF2 protein as a platform for metal ion detection. Few examples of GAF or other bilin-binding proteins have been tested as metal ion sensors or used in multi-protein sensing platforms.^26,38–40^ NIR bacteriophytochrome proteins have been used to develop Ca^2+^ and Hg^2+^ sensors.^38,39^ Some bacteriophytochrome-derived miRFPs can also be inherently quenched by Cu^2+^ ions with picomolar dissociation constants.^40^ The PEB-bound GAF domain from *Spirulina subsalsa* (NCBI WP_017306776.1) was also reported to be quenched by Cu^2+^ ions with apparent *K*_D_ of ∼15 µM.^26^ Although free bilins in solution can bind metal ions, as can unfolded bilin-binding proteins, no examples where metal ions bind directly to bilins embedded within phytochrome proteins have been reported.^41–43^

Here we investigated the suitability of All1280 GAF2-PEB as a metal sensor. We show that GAF2-PEB undergoes a significant and reversible zinc-induced fluorescence enhancement with pH-dependent dissociation constants ranging from low micromolar to ∼20 nM. UV-visible absorption data is consistent with direct binding of zinc to the protein-bound bilin cofactor. Sterically modified site-directed mutants retain or show reduced Zn^2+^-induced fluorescence. Conversely, Slr1393 GAF3-PEB fluorescence was not increased with Zn^2+^ and GAF3 does not harbor a DXCF motif cysteine residue near the bilin cofactor. Mutation of GAF2 to replace this cysteine residue with alanine led to a mutant that did not show fluorescence enhancement or any changes to the UV-visible absorption spectrum with Zn^2+^ ions, confirming that the DXCF cysteine residue is required for Zn^2+^-induced fluorescence. Given the small size, high quantum yield, and substantial zinc-induced fluorescence response, GAF2-PEB is an excellent candidate for developing higher affinity metal-sensing scaffolds and provides insight to how bilin-binding fluorescent proteins can be used to develop new metal-sensing platforms.

## Materials and Methods

### General Information

All reagents were purchased from commercial sources and used as received. Primers were ordered from Sigma-Aldrich. *E. coli* strain (DH5α) for cloning and *E. coli* strain BL21(DE3) for protein expression were obtained from New England Biolabs. Restriction enzymes, T4 DNA ligase, Q5 DNA polymerase, and DNase I were also obtained from New England Biolabs. PCR cleanup, plasmid DNA miniprep, and gel extraction kits were from New England Biolabs. Columns for purification were from Bio-Rad. Protein gel electrophoresis was carried out using Bio-Rad 4–20% Mini-PROTEAN TGX Stain-Free Protein Gels. Gels were imaged on a Bio-Rad ChemiDoc Touch Imaging System. Fast Protein Liquid Chromatography was carried out on a Bio-Rad NGC Quest 10 Chromatography system. UV-visible absorption spectra were collected on an Agilent Cary 100 UV-visible spectrophotometer and fluorescence spectra on an Agilent Eclipse spectrofluorimeter using quartz cuvettes (Starna cells) with 1 cm path lengths. All measurements were conducted at 25.0 °C, maintained by a circulating water bath. To remove adventitious metal ions, all buffers used for spectroscopy were treated with Chelex-100 resin (Bio-Rad) according to the manufacturer’s batch protocol. Experiments involving Cu^+^ were done anaerobically using buffer that was freshly degassed with argon and freshly prepared [Cu(CH_3_CN)_4_]PF_6_ stock solutions (in acetonitrile). Anaerobic manipulations were done in an anaerobic chamber or using cuvettes with a screw cap and septum and a gas-tight Hamilton syringe.

### Molecular Cloning

The plasmid pSR43.6r (Addgene no. 63197) encoding heme oxygenase (HO1, UniProtKB P72849) and phycocyanobilin:ferredoxin oxidoreductase (PcyA, UniProtKB Q55891) was a gift from Jeffrey Tabor.^44^ The plasmid pACYC-PebS-HO1 was constructed in pACYCDeut-1 (Novagen) using the HO1 gene from pSR43.6r and the PebS gene from *Prochlorococcus phage P-SSM2* (UniProtKB Q58MU6, codon optimized for *E. coli*). An additional T7 terminator and T7 promoter were placed between the PebS and HO1 genes to facilitate multi-protein expression. The pET28-GAF3 plasmid (ordered from Genscript) was constructed using the gene Slr1393 of *Synechocystis* PCC 6803 (NCBI protein ID BAA17210, UniProtKB P73184, amino acids L441–Q596) with a TGA stop codon and inserted between BamHI and XhoI restriction sites in pET28a. All plasmids were verified by DNA sequencing.

*pET28a-GAF2*. The gene encoding for the GAF2 domain from All1280 of *Nostoc* PCC 7120 (NCBI protein ID BAB73237.1, UniProtKB Q8YXD3, amino acids 562-727) was ordered from Genscript (codon optimized for *E. coli*) in the pUC18 cloning vector. GAF2 was PCR amplified from pUC18-GAF2 using the primers listed in Table S1 and the PCR product with a TAA stop codon was inserted between BamHI and XhoI restriction sites in pET28a-GAF3. The nucleotide and amino acid sequences of the coding region of the resulting plasmid, pET28a-GAF2, are shown in Figure S1.

*pET28a-GAF2*_*L652A*_, *pET28a-GAF2*_*L652I*_, *pET28a-GAF2*_*L652W*_, *and pET28a-GAF2*_*C620A*_. The plasmids encoding for GAF2 mutants were prepared by Q5 site-directed PCR mutagenesis (NEB) of pET28a-GAF2 using the primers listed in Table S1.

### Expression and Purification of GAF2 and GAF3 Proteins

The pACYC-ho1-pebS and pET28a-GAF2 (or GAF2 mutants) or pACYC-ho1-pebS and pET28a-GAF3 plasmids were co-transformed into *E coli* BL21(DE3) cells and grown on LB agar plates with kanamycin (50 µg/mL) and chloramphenicol (25 µg/mL). Single colonies were selected and starter cultures were grown overnight at 37 °C in terrific broth (TB), as previously reported.^45^ Protein expression starter cultures were inoculated (1:100) into unbaffled flasks. The cultures were grown at 37 °C and 200 rpm to OD_600_ of 0.6-0.7. Protein expression was induced with isopropyl beta-*D*-1-thiogalactopyranoside (IPTG, 100 µM). The temperature was lowered to 18 °C after induction and the cells were grown for an additional 18 hours with shaking speed reduced to 120 rpm. The cells were centrifuged at 5000 ’
s *g* for 10 minutes at 4 °C and pelleted. The cell paste was resuspended in 25 mL phosphate buffer (20 mM phosphate buffer, 200 mM NaCl, pH 8) containing phenylmethanesulfonyl fluoride (PMSF, 1mM), DNase I, CaCl_2_ (0.5 mM), MgCl_2_ (10 mM) and one EDTA-free Pierce Protease Inhibitor tablet. The resuspended cell paste was sonicated on ice and the lysed cells were centrifuged at 21000 ’ *g*, 4 °C for 1 hour. The supernatant containing the protein was filtered (0.45 µm) and applied on an IMAC Ni^2+^-NTA column (Bio-Rad) for FPLC purification. The column was washed with five column volumes of wash buffer (20 mM phosphate buffer, 200 mM NaCl, pH 8) and the protein eluted with 2 column volumes of elution buffer (wash buffer including 500 mM imidazole, pH 8). The appropriate fractions were pooled and transferred into metal-free HEPES buffer (50 mM HEPES, 100 mM NaCl, pH 7.1) using either a 10,000 molecular weight cut-off filter (Millipore) or through a Mini Bio-Gel P-6 Desalting Cartridge (Bio-Rad). The concentration of the protein was determined by using the absorption measured at 280 nm and the corresponding extinction coefficient of 20970 M^-1^cm^-1^.

Thrombin protease (Cytiva) was used to remove the histidine tag for some experimental analyses. The (His_6_)GAF protein sample was treated with thrombin protease (1 unit/mg protein) for 16 hours at 4 °C. The reaction was stopped by the addition of PMSF (1 mM) and the mixture purified using the IMAC Ni^2+^-charged column. The histidine-free GAF fractions were pooled and transferred into metal-free HEPES buffer (50 mM HEPES, 100 mM NaCl, pH 7.1) using either a 10,000 molecular weight cut-off filter (Millipore) or through a Mini Bio-Gel P-6 Desalting Cartridge (Bio-Rad).

### Spectroscopic Measurements

For all spectroscopic measurements, a final concentration of 3–14 µM purified protein was prepared in 2.5 mL of Chelex-treated buffer (50 mM HEPES, 100 mM NaCl, pH 7.1 except where indicated). Fluorescence data were obtained by exciting at 495 nm and collecting the emission from 505–800 nm. Fluorescence quantum yields for the GAF2-PEB mutants were measured using the known Φ = 0.31 for GAF3-PEB as a standard.^25^

### Metal Ion Sensitivity Assays

The effects of metal ions on the fluorescence of each protein were screened by comparing the fluorescence of 3-4 µM protein sample in buffer (50 mM HEPES, 100 mM NaCl, pH 7.1) before and after the addition of KCl, MgCl_2_, or CaCl_2_ (800 µM each), and ZnCl_2_, CoCl_2_, MnCl_2_, NiCl_2_, CuCl_2_, FeCl_2_, or [Cu(CH_3_CN)_4_]PF_6_ (100 µM, Fe^2+^ and Cu^+^ stocks were prepared fresh and used immediately). The [Cu(CH_3_CN)_4_]PF_6_ stock solution was prepared in acetonitrile and all other metal stock solutions were prepared using mQ water. Ten minutes of mixing time was allowed to ensure equilibration. For zinc and copper experiments, EDTA (500 µM, for Zn^2+^ and Cu^2+^) or DTT (200 µM, for Cu^+^) was added to test the reversibility of the metal-induced fluorescence response. Thirty minutes of mixing time was allowed to ensure equilibration.

### UV-visible Absorption Analysis of GAF2-PEB and GAF2_C620A_-PEB with Zinc

The UV-visible absorption spectra for GAF2-PEB (or GAF2_C620A_-PEB) and GAF2-PEB (or GAF2_C620A_-PEB) with Zn^2+^ and Zn^2+^ + EDTA were collected in buffer (50 mM HEPES, 100 mM NaCl, 1 mM ATP). After collecting the initial absorption spectrum of GAF2-PEB (or GAF2_C620A_-PEB, 14 µM), ZnCl_2_ (250 µM) was added and allowed to equilibrate up to 10 min, and the spectrum collected. EDTA (1 mM) was subsequently added, allowed to equilibrate for up to 1 h, and the absorption spectrum collected.

### Zinc and Copper Titrations

All dissociation constants were determined at pH 7.1 (except as indicated for GAF2-PEB below) using the change in fluorescence emission. Zn^2+^ titrations were carried out with ADA (1 mM, GAF2-PEB, (His_6_)GAF2-PEB, (His_6_)GAF3-PEB, or (His_6_)GAF2_L652I_-PEB) or ATP (1 mM, GAF2-PEB) to buffer Zn^2+^ concentrations. All Cu^2+^ titrations were carried out with NTA (1 mM). GAF2-PEB titrations with Zn^2+^ were also carried out at pH 6.5, 7.0, 7.5, and 8.0. Chelex-treated buffers used for each pH are as follows: 50 mM MES, 100 mM NaCl for pH 6.5; 50 mM HEPES, 100 mM NaCl for pH 7.0-8.0. Free metal ion concentrations were calculated with MaxChelator.^46^ The dissociation constants used by MaxChelator for Zn^2+^ binding to ADA and ATP are as follows: 9.40 × 10^−6^ M (ATP, pH 7.0); 8.90 × 10^−6^ M (ATP, pH 7.1); 1.49×10^−5^ M (ATP, pH 6.5); 8.78×10^−8^ M (ADA, pH 7.1); 7.17×10^−8^ M (ADA, pH 7.5); 6.44×10^−8^ M (ADA, pH 8.0).^46^ The dissociation constant for Cu^2+^ binding to NTA at pH 7.1 is 5.30×10^−11^

M.^46^ For each titration, aliquots of ZnCl_2_ or CuCl_2_ were added to the protein (3-4 µM). With each addition, the fluorescence emission spectrum was recorded after mixing for 5 minutes (no further changes to the spectra were observed beyond this time). The integrated fluorescence intensity (505–800 nm, normalized to the initial protein fluorescence) was plotted against the total metal concentration for each addition. The resulting binding isotherms for Zn^2+^ were fitted to single site or two site models (accounting for ADA or ATP) for apparent dissociation constants, *K*_D_, using DynaFit (http://www.biokin.com/dynafit/). Figure S2 shows example fitting scripts for each type of binding model. Titrations with Cu^2+^ were not fitted for dissociation constants given the lack of reversibility in Cu^2+^ binding and quenching.

### Assessment of pH-dependent Fluorescence Response of GAF2-PEB, GAF2_L652I_-PEB, and GAF2_C620A_-PEB

The fluorescence of the PEB-bound protein initially, with ZnCl_2_, and with ZnCl_2_ + EDTA was measured at pH values ranging from 5.5 to 9.0 (50 mM MES, 100 mM NaCl for pH 5.5-6.5; 50 mM HEPES, 100 mM NaCl for pH 7.0-8.0; 50 mM CHES, 100 mM NaCl for pH 8.5-9.0). For each pH, a sample of GAF2-PEB (or mutant, 3-4 µM) was added to buffer in a quartz cuvette and the fluorescence spectrum was recorded. Excess ZnCl_2_ (100 µM) was added and mixed for up to 10 min, then the fluorescence spectrum collected. Finally, EDTA (500 µM) was added and after equilibration for 30 min, the final fluorescence spectrum was collected.

## Results and Discussion

### Protein Expression and Characterization of GAF2-PEB, GAF3-PEB, and GAF2-PEB Mutants

GAF proteins were expressed by co-transforming a plasmid encoding for the GAF protein and a plasmid encoding heme oxygenase and PebS enzymes into competent *E. coli* cells. Heme oxygenase and PebS enzymes catalyze the synthesis of phycoerythrobilin (PEB) from cellular heme,^27,30^ which then autocatalytically reacts with the overexpressed GAF domain to form a covalent thioether bond with a conserved cysteine residue (Figure 1A, Scheme S1). GAF2-PEB can also autoisomerize to GAF2-PUB, resulting in a mixture of species that can be detected by UV-visible absorption spectroscopy (Figure 1A).^25^ The N-terminal His_6_ tag on each GAF protein was used for Ni^2+^-NTA affinity chromatography purification. For comparison of results in the presence and absence of a His_6_ tag, the His_6_ tag was cleaved from GAF2-PEB using thrombin protease. Protein purity was confirmed by denaturing gel electrophoresis (Figure S3). The spectroscopic properties of GAF2-PEB and GAF3-PEB were similar to those previously reported (Figure 1B-C, S4, Table S2). The spectroscopic properties of the GAF2-PEB mutants will be discussed below (Figure S4).

**Figure 1.**
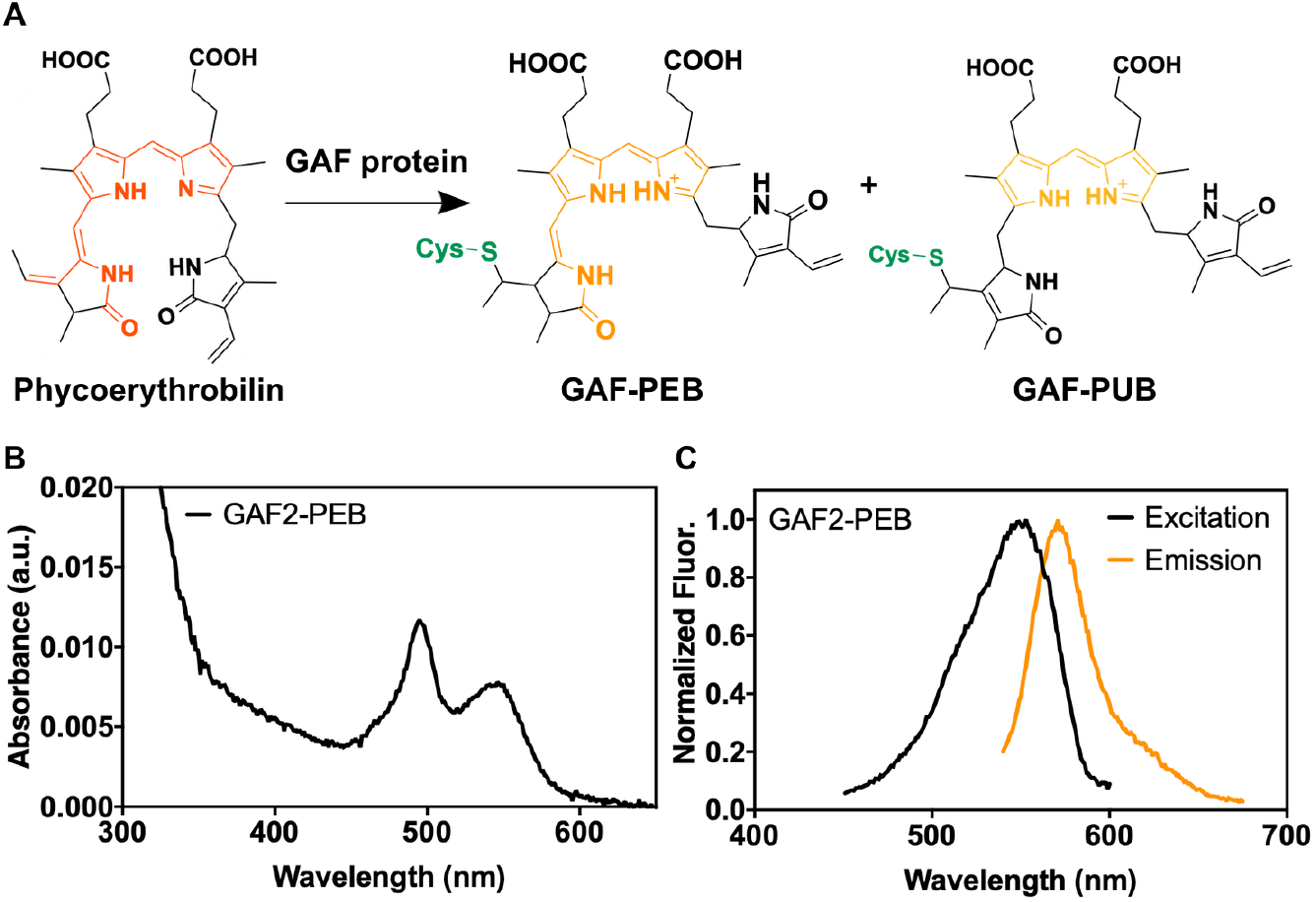
Chemical structure and spectroscopic properties of phycoerythrobilin (PEB)-bound GAF proteins. A) General chemical structure of PEB in solution and upon reaction with a GAF protein cysteine residue. Partial autoisomerization of GAF-PEB to phycourobilin (PUB) can occur upon bilin binding.^25^ B) UV-visible and C) fluorescence excitation (λ_em_ = 620 nm) and emission (λ_ex_ = 495 nm) spectra of purified GAF2-PEB (10 µM in B and 3 µM in C). Buffer: 50 mM HEPES, 100 mM NaCl, pH 7.1.

### Metal Sensitivity of GAF2-PEB and GAF3-PEB Proteins

To determine whether the fluorescence of cyanobacteriochrome GAF-PEB proteins could be affected by the presence of metal ions, we measured the fluorescence of GAF2-PEB and GAF3-PEB in the presence and absence of a series of several biologically relevant metals (Figure 2A-B). Previous work on a related GAF protein from *Spirulina subsalsa* showed that excess Cu^2+^ quenched the fluorescence by ∼75%, whereas no effect in the presence of some other biologically relevant metals like Zn^2+^, Ni^2+^, Mn^2+^, and Co^2+^ was observed.^26^ Here, we found that GAF2-PEB was similarly quenched by Cu^2+^ by ∼66% and GAF3 was quenched by ∼61% (Figure 2A-B, S5). We also tested Cu^+^, given that this is the redox state expected to predominate within the reducing environment of live cells, and found that ∼66% of the fluorescence of GAF2-PEB was quenched and ∼70% of the fluorescence of GAF3-PEB was quenched (Figure 2, S5). More interestingly, GAF2-PEB showed a significant increase in fluorescence in the presence of Zn^2+^ that was not observed for GAF3-PEB at physiologically relevant zinc concentrations (Figure 2A-B, S6). A slight Zn^2+^-induced fluorescence enhancement was observed for GAF3-PEB with excess zinc (Figure 2B, S5). Similarly, no effects from Zn^2+^ on the fluorescence of the *S. subsalsa*-derived GAF-PEB were reported.^26^ Examination of the fluorescence spectra in the presence and absence of Zn^2+^ reveals substantial changes in the fluorescence spectrum for GAF2-PEB (Figure 2C). Upon binding Zn^2+^, the fluorescence emission maximum for GAF2-PEB shifts from 572 nm to 517 nm. At the same time, a low-intensity broad band in the red-shifted region appears at ∼593 nm. The initial maximum is also obscured by another shoulder band centered around 547 nm. We also tested whether the significant effects induced by Cu^+/2+^ and Zn^2+^ on GAF2-PEB were reversible, and found that while the quenching induced by Cu^2+^ was not reversible and by Cu^+^ was only partially reversible, the zinc-induced fluorescence enhancement could be completely reversed with addition of EDTA (Figure 2C-E). For GAF3-PEB, we found that Cu^+/2+^-induced fluorescence quenching were mostly irreversible, and the effects induced by excess Zn^2+^ were partially reversible (Figure S5).

**Figure 2.**
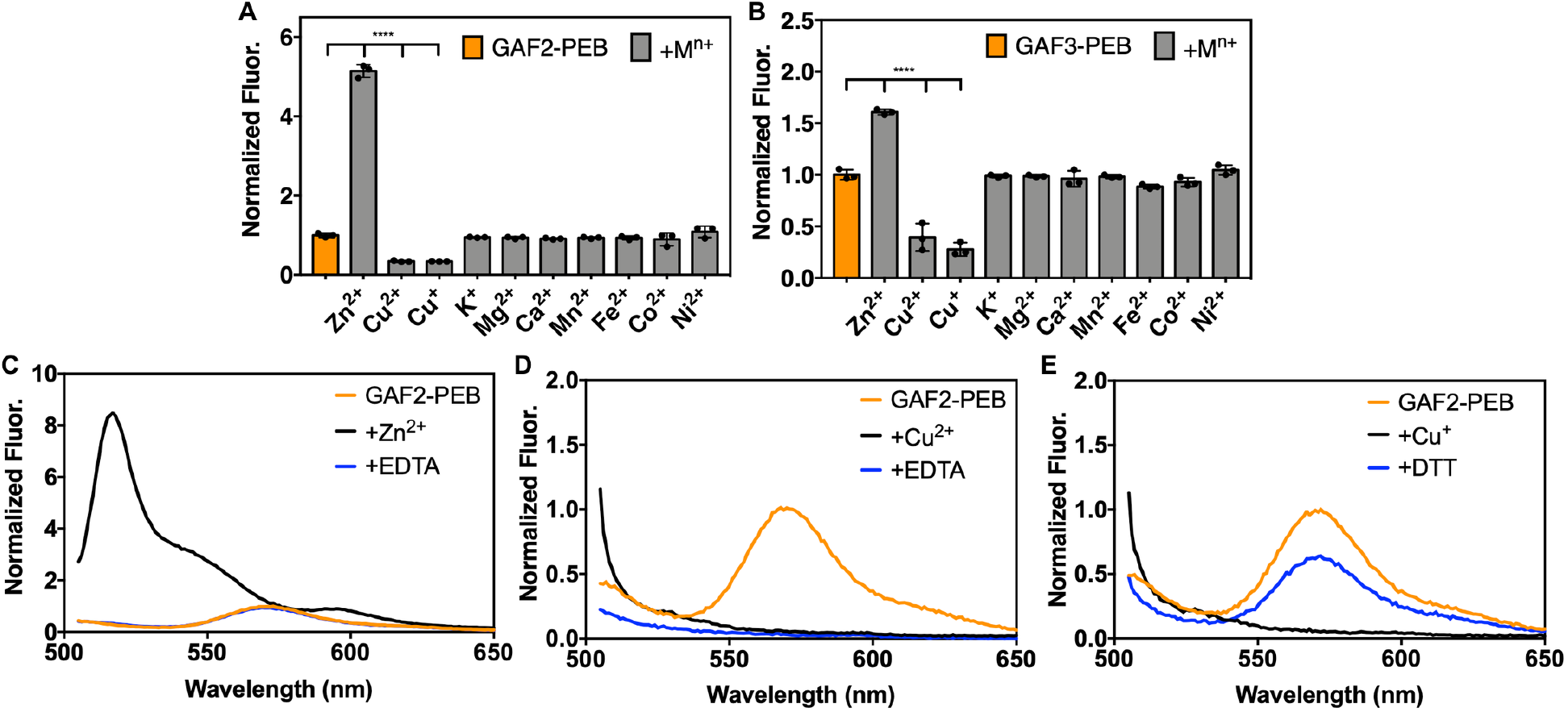
Metal sensitivity and reversibility of GAF2-PEB and GAF3-PEB. A-B) To 3 μM of protein, 266 eq (800 µM K^+^, Mg^2+^, or Ca^2+^) or 33 eq (100 µM Zn^2+^, Cu^2+^, Cu^+^, Mn^2+^, Fe^2+^, Co^2+^, or Ni^2+^) metal salt was added. Data are shown as the integrated fluorescence normalized to the initial integrated fluorescence of the protein. λ_ex_ = 495 nm and λ_em_ = 505-800 nm. Data were analyzed by means of one-way analysis of variance; ****p < 0.0001. Individual data points are overlaid on the bar chart representation. All error bars represent the standard deviation for three replicates. C) Fluorescence emission spectra of (His_6_)GAF2-PEB (3 µM) initially (orange line), upon addition of ZnCl_2_ (100 µM, black line), and with addition of EDTA (500 µM, blue line). D) Fluorescence emission spectra of (His_6_)GAF2-PEB (3 µM) initially (orange), upon addition of CuCl_2_ (100 µM, black), and with addition of EDTA (500 µM, blue). E) Fluorescence emission spectra of (His_6_)GAF2-PEB (3 µM) initially (orange), upon addition of [Cu(CH_3_CN)_4_]PF_6_ (100 µM, black), and with addition of DTT (200 µM, blue). Fluorescence spectra were normalized to the maximal emission intensity of GAF2-PEB at 572 nm. Buffer: 50 mM HEPES, 100 mM NaCl, pH 7.1.

The fluorescence response of GAF2-PEB to Zn^2+^ is unique among folded bilin-binding fluorescent proteins, although a Zn^2+^-induced fluorescent enhancement was reported for a GFP mutant harboring a tridentate half-porphyrin-like chromophore.^47^ Free bilins like BV can coordinate via the pyrrole rings to Zn^2+^ and Cu^2+^ ions to form stable complexes.^42^ Binding of Zn^2+^ can lead to increased fluorescence due to a decrease in flexibility of the chromophore, and is a feature that has been exploited to detect covalent bilin binding to these proteins.^41,48^ For this method, bilin-binding proteins are subjected to denaturing gel electrophoresis in the presence of a Zn^2+^ “stain” and only proteins harboring a covalently bound bilin cofactor will fluoresce red on the gel. For example, while the folded GAF-PEB protein from *S. subsalsa* does not undergo any changes in fluorescence in the presence of Zn^2+^, the denatured protein fluoresces.^26^ To determine whether Zn^2+^ may be binding directly to the bound PEB cofactor embedded within GAF2, we compared the UV-visible absorption spectra obtained in the presence and absence of zinc (Figure 3). We expect that metal binding to the bilin cofactor would change the absorption spectrum, as observed for some metals that bind free bilins, including Zn^2+^ binding to free PEB.^42,43^ Here we found that the absorption maxima at 497 nm and 542 nm shifted to 510 nm with a shoulder at ∼540 nm and 587 nm upon the addition of zinc at pH 7.1, suggesting direct binding to the chromophore. The red-shifting of these peaks is consistent with the absorption spectra reported for Zn^2+^ binding to isolated PEB and to phycoerythrin proteins.^43,49^

**Figure 3.**
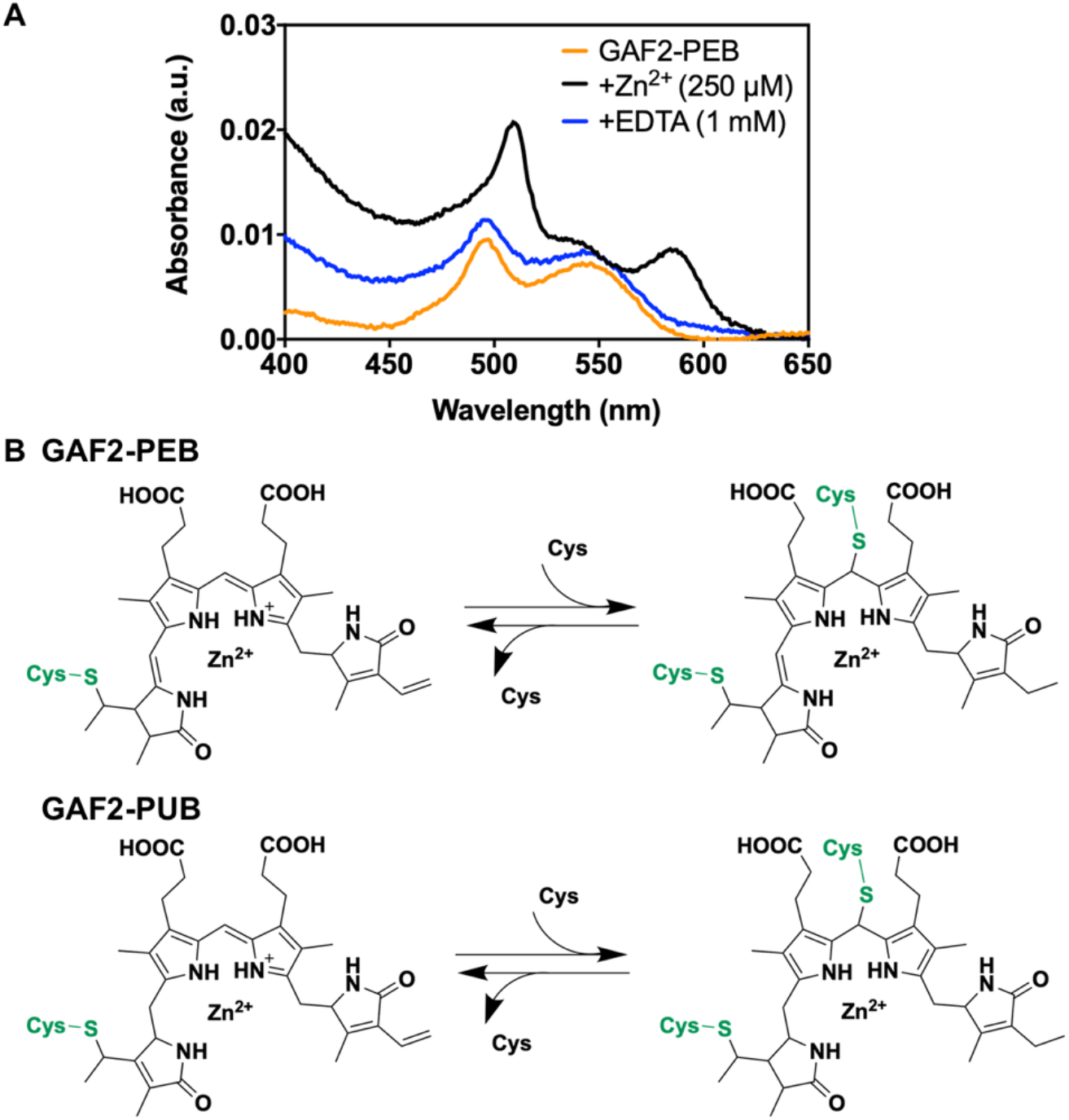
A) UV-visible absorption spectra GAF2-PEB (14 µM, orange line) initially, with addition of ZnCl_2_ (250 µM, black line), and with addition of EDTA (1 mM, blue line). Buffer: 50 mM HEPES, 100 mM NaCl, 1.0 mM ATP, pH 7.1. B) Chemical structure schematic depicting the possible GAF2-bilin structures present upon binding PEB and proposed location for zinc binding. Briefly, covalently bound PEB can be present in GAF2 bound to one or two cysteine residues (with the second cysteine from the DXCF domain). GAF2-PEB can also partially autoisomerize to GAF2-PUB,^25^ which could also bind to one or two cysteine residues, although the latter has not been experimentally demonstrated and is not expected to be fluorescent due to a lack of extended conjugation.

### Copper and Zinc Titrations for GAF2-PEB

To determine the affinity of GAF2-PEB to zinc, we titrated the protein with increasing amounts of zinc in the presence of adenosine triphosphate (ATP) as a buffering chelator to control the available Zn^2+^ concentrations. Plotting the integrated fluorescence for each spectrum against the total zinc concentration led to a binding isotherm with increasing fluorescence that could be fitted to a single site binding model with *K*_D_ = 1.1 ± 0.1 µM (Figure 4, Table S3). Similar titrations were carried out in the presence of *N*-(2-acetamido)iminodiacetic acid (ADA) and although the fluorescence enhancement did not saturate, the data could be fitted to yield a similar *K*_D_ of 0.65 ± 0.01 µM (Figure S7, Table S3). The presence of a His_6_ tag on GAF2-PEB yielded only a slightly stronger *K*_D_ of 0.52 ± 0.01 µM under the same conditions, suggesting that the His_6_ tag does not impact the Zn^2+^-induced fluorescence response. Such a micromolar dissociation constant is on the weaker side for physiological zinc concentrations that are typically picomolar to nanomolar in many cells, including bacteria, but could fall in the range of zinc concentrations found in specialized cells, upon application of extracellular stimuli, or with growth in high zinc medium.^36,50,51^ We also titrated GAF2-PEB with Cu^2+^ in the presence of nitrilotriacetic acid (NTA), but given the lack of reversibility in Cu^2+^-induced quenching it is not feasible to fit this data to a binding model (Figure S8). There may be multiple possible mechanisms for Cu^2+^-induced quenching, both in the presence and the absence of a His_6_ tag. We believe that it is possible for copper to bind directly to the cofactor and induce fluorescence quenching, but copper may also bind to the protein and induce quenching through other energy or electron transfer-based quenching mechanisms, as described in prior work on BV-binding miRFPs.^40^

**Figure 4.**
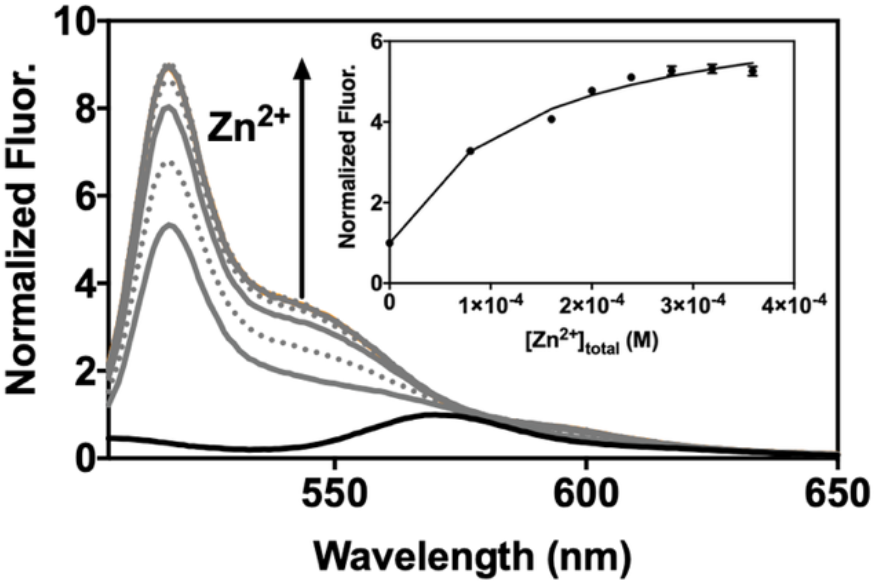
Zn^2+^ binding titration for GAF2-PEB at pH 7.0. Normalized fluorescence spectra of GAF2-PEB (3 µM) titrated with increasing amounts of ZnCl_2_ in the presence of ATP (1 mM). Inset: normalized integrated fluorescence plotted against total Zn^2+^ concentration. Fluorescence was normalized relative to the initial GAF2-PEB fluorescence. Data were fitted with DynaFit (Experimental Section, Figure S2). λ_ex_ = 495 nm, λ_em_ = 505-800 nm. Buffer: 50 mM HEPES, 100 mM NaCl, pH 7.0. All error bars represent the standard deviation for three replicates.

For comparison, we titrated GAF3-PEB with Zn^2+^ and Cu^2+^. Here, we observed fluorescence quenching for both metals over the concentration range of the titrations (Figure S7, S8). For Zn^2+^, ADA was used as the buffering chelator, and the resulting binding isotherm could be fitted to a single site binding model to yield a *K*_D_ of 44 ± 4 nM (Table S3). This *K*_D_ supports binding of Zn^2+^ to GAF3-PEB that is stronger than that observed for Zn^2+^ binding to GAF2-PEB at pH 7.1, but given the quenching response rather than a fluorescence enhancement, it likely originates from a different mechanism and probably a different binding site. Similarly, titration with Cu^2+^ induced quenching. As for GAF2-PEB, Cu^2+^-induced quenching is not reversible and therefore it is not feasible to fit this data to a binding model (Figure S8).

### Effect of pH on Zn^2+^-Induced Fluorescence Enhancement of GAF2-PEB

To determine the potential of using GAF2-PEB as a zinc sensor in biological systems, we tested the effect of pH on the Zn^2+^-induced fluorescence enhancement. From pH 6.0 to 9.0, significant increases in the Zn^2+^-induced fluorescence over the initial GAF2-PEB fluorescence were observed and range from ∼3-fold to 7-fold from pH 6.0-7.5 (Figure 5A). Very little enhancement was observed with addition of Zn^2+^ at pH 5.5. At higher pH, the initial fluorescence decreases significantly, yet addition of Zn^2+^ leads to a similar integrated fluorescence level, so the overall enhancement ranges from ∼10-20-fold from pH 8.0 to 9.0 (Figure 5A-C). The level of reversibility decreases with increasing pH whereby the response is mostly reversible up to pH 7.0, partially reversible at pH 7.5 and 8.0, and mostly irreversible from pH 8.5 to 9.0 (Figure 5A,D). The lack of fluorescence enhancement at pH 5.5 suggests that the metal binding residues are still protonated. From pH 6.0 some deprotonation has occurred, allowing the zinc binding site to form. Examination of the fluorescence spectrum with further increasing pH reveals a probable change in the fluorophore environment at higher pH, given the loss of the shoulder band at 547 nm and the loss of reversibility upon addition of chelator. We hypothesize that at these higher pH values, deprotonation of the PEB cofactor could occur, leading to both a loss in the initial fluorescence and, if involved in zinc binding, an increase in binding affinity. To test this hypothesis, we performed zinc binding titrations at pH 6.5, 7.5, and 8.0 to compare with the titration at pH 7.0 (Figure 4). At pH 6.5 and in the presence of ATP, titration of GAF2-PEB Zn^2+^ led to a binding isotherm that could be fitted to a single site binding model, similar to pH 7.0, but with a slightly weaker dissociation constant of 5 ± 1 µM (Figure S9A, Table S3, compared to 1.1 µM at pH 7.0). A similar titration at pH 8.0 could not be fitted to a single site model, but fitting to a two-site model suggests stronger dissociation constants at ∼20 nM and ∼80 nM (Figure S9, Table S3). Titration at pH 7.5 could not be fitted well to either a single or two site model (Figure S9), suggesting there could be a mixture of species present at this intermediate pH. Collectively, these results support the utility of GAF2-PEB as a Zn^2+^ sensor over a biologically relevant range from pH 6.0-8.0, and suggest that Zn^2+^ binding might be modulated via mutations that alter the metal binding site p*K*_a_ and affinity.

**Figure 5.**
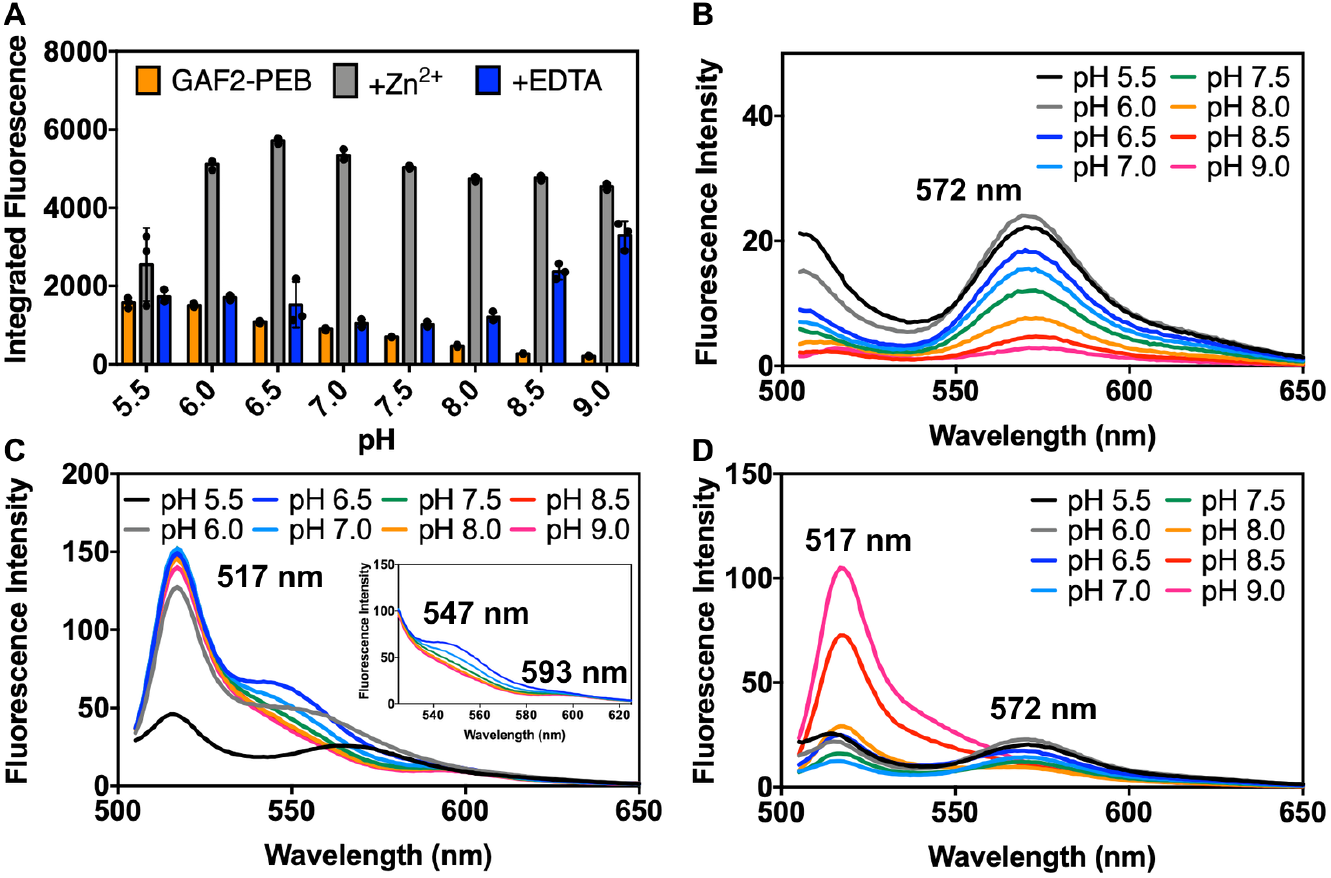
Effect of pH on the fluorescence intensity of GAF2-PEB in the presence and absence of Zn^2+^. A) Integrated fluorescence intensity of GAF2-PEB (orange, 3 μM), GAF2-PEB with the addition of ZnCl_2_ (grey, 100 μM), and GAF2-PEB + Zn^2+^ with the addition of EDTA (blue, 500 µM). λ_ex_ = 495 nm and λ_em_ = 505-800 nm. Individual data points are overlaid on the bar chart representation. All error bars represent standard deviation for 3 replicates. B) Spectra for GAF2-PEB representing the data shown in part A. C) Spectra for GAF2-PEB + Zn^2+^ representing the data shown in part A. Inset shows an expansion of the region from 525-625 nm. D) Spectra for GAF2-PEB + Zn^2+^ + EDTA representing the data shown in part A. Buffers: 50 mM MES, 100 mM NaCl (pH 5.5, 6.0, 6.5); 50 mM HEPES, 100 mM NaCl (pH 7.0, 7.5, 8.0); 50 mM CHES, 100 mM NaCl (pH 8.5, 9.0).

### Comparison of GAF2 and GAF3 Sequences and Structures

Given that the fluorescence and absorption spectra for Zn^2+^ and GAF2-PEB suggest that zinc could bind directly to the PEB chromophore to induce fluorescence enhancement, and that there are comparatively minimal effects of Zn^2+^ on GAF3-PEB, we compared the sequences and structures of GAF2 and GAF3. The X-ray crystal structure of GAF3-PCB was previously reported, but the structure of GAF2 has not been published.^52^ Using Clustal Omega,^53^ the sequences of GAF2 and GAF3 were aligned with ∼40% sequence identity (Figure 6A). We used SwissModel to generate a homology model of GAF2, which we compared with the structure of GAF3 (Figure 6B).^54^ The bilin cofactor was modeled by aligning PCB-bound GAF3 with the homology model of GAF2. PCB is similar to PEB with the exception of the lack of the Δ15, 16 double bond (Scheme S1). The homology model of GAF2 was based off of a similar GAF domain, which showed the highest sequence identity (49.67%) among the template options and resulted in good QMeanDisCo and GMQE scores (0.75 ± 0.05 and 0.76, respectively), representing the model quality in coverage-dependent (GMQE) and non-coverage dependent (QMeanDisCo) ways.^54,55^ A similar model with the same scores was obtained using PDB 6OAQ (Figure S10). Several other templates including GAF3 (PDB 5DFX) were tested as models and the corresponding scores are presented in Table S4. By comparing the surface rendering of the GAF2 6OAP model with the surface rendering of the crystal structure of GAF3, which overlay well, we found an apparent increase in solvent access to the chromophore for GAF2 (Figure 6B-D). A similar solvent access area in GAF2 was identified in the other top homology model based on 6OAQ (Figure S10). Much of the access to the bilin cofactor in GAF2 could be due to decreased packing of the loop region below the cofactor in the GAF2 homology model. Given the flexibility of the loop region, we compared several other templates as models and found that in most, the bilin-binding pocket remained more accessible than in GAF3, although it was heavily affected by the predicted folding of the loop region (Figure S11). Using the highest-scoring models based on 6OAP and 6OAQ, we identified a residue (Leu652) on a more structurally-stabilized α-helical region above the binding pocket that could affect the steric access to the bilin cofactor. Even in the lower-scoring models with a more packed loop region, the Leu652 residue remained well positioned to affect solvent access to the chromophore. In GAF3, the corresponding residue is Asn. We hypothesized that mutation of this amino acid could allow control of solvent and thereby metal access to the bilin and be used to control the metal-induced fluorescence response. To test the hypothesis that the Leu652 residue could affect metal access to the PEB cofactor, we prepared two single site mutants of GAF2 where Leu was replaced by Ala or Ile. The Ala mutation would increase the solvent access to the chromophore whereas the Ile mutation would retain hydrophobicity but potentially decrease solvent access to the chromophore. A third single site mutant in which Leu652 was replaced by tryptophan was not chromophorylated during protein expression and was therefore not further characterized (data not shown).

**Figure 6.**
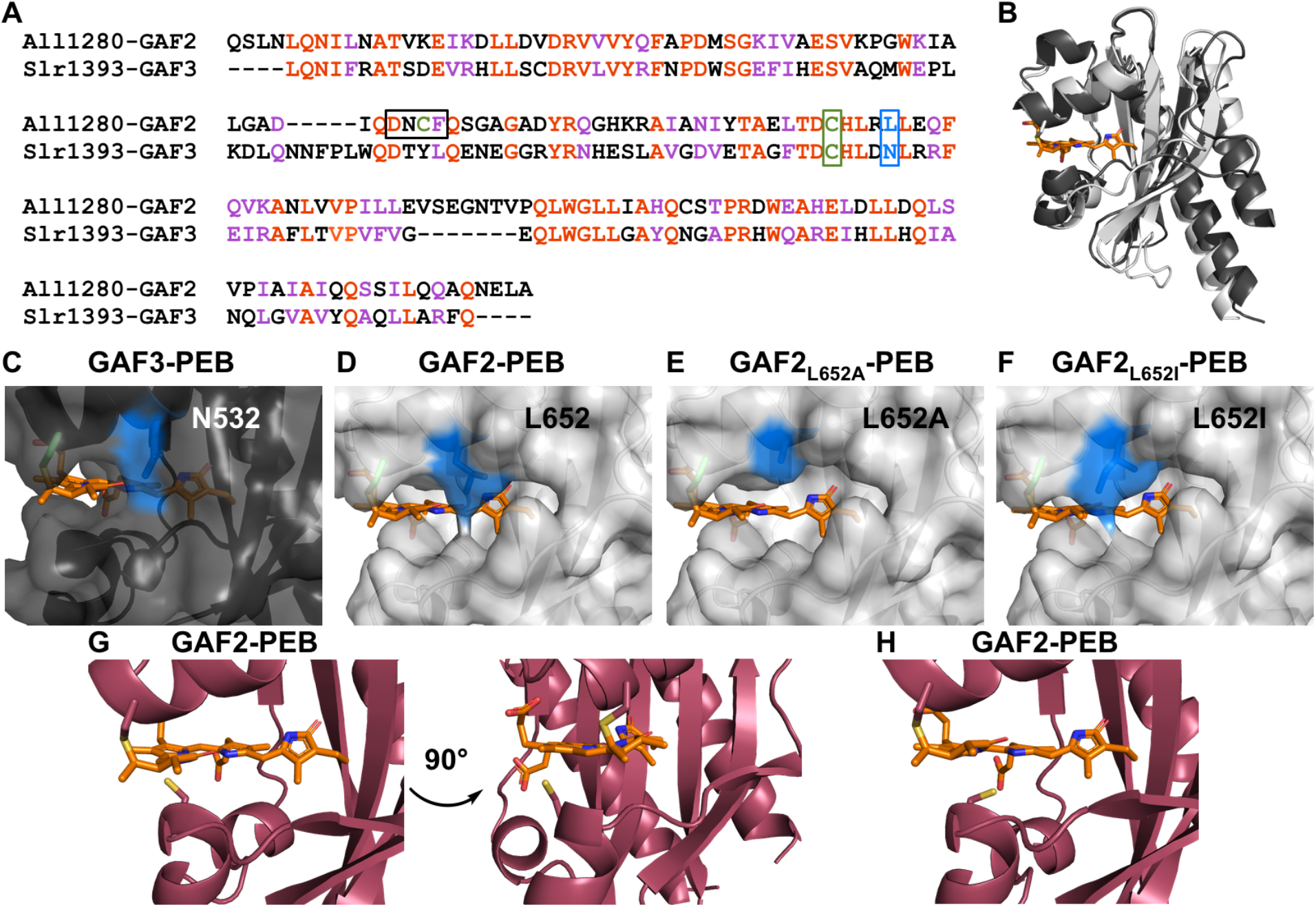
Comparison of GAF proteins. A) Sequence alignment of All1280 GAF2 and Slr1393 GAF3. Amino acids are #441-596 from Slr1393 and #526-727 from All1280. The aligned residues are colored red and similar residues are colored purple. One of the residues mutated in GAF2 in this work (L652) is colored blue and outlined and the conserved cysteine residue (C648) that binds to the bilin in both GAFs is colored green and outlined. The corresponding residues are N532 and C528 in GAF3. The DXCF cysteine residue 620 found only in GAF2 is also highlighted green and the whole DXCF motif is outlined in the sequence alignment. B) Overlay of PyMOL representations of the crystal structure of Slr1393 GAF3-PCB (black, PDB 5DFX) and homology model of All1280 GAF2 (grey, prepared using SwissModel and based on PDB 6OAP). C-F) PyMOL models of the PEB binding site in C) GAF3, D) GAF2, E) GAF2_L652A_, and F) GAF2_L652I_. Mutations were modeled using the PyMOL mutagenesis wizard. The N532 side chain in GAF3 (C) and side chains in position 652 of GAF2 (D-F) are colored blue. G) Two views of the PyMOL representation of the PEB binding site in GAF2 (maroon, prepared using SwissModel and based on PDB 6MGH) shown with the nearby DXCF C620 residue. H) Depiction of an additional possible rotamer for C620 from part G that could place this residue even closer to the center of the bound bilin cofactor.

In addition to the possible role of sterics around the proposed zinc binding site, we also considered whether the DXCF motif, found in GAF2 but not GAF3, plays a role in Zn^2+^-induced fluorescence enhancement. Here, we examined the location of Cys620 in the homology models prepared for GAF2 as described above. Among the six models, three placed the Cys620 in the same position, oriented within ∼3-4 Å of the C10 position on the bilin cofactor (Figure 6G and S12, models based on PDB 6MGH, 5DFX, and 7LSC). This position also places Cys620 and another rotamer of Cys620 within a Zn^2+^-S_Cys_ binding distance range of ∼2.5-3.5 Å (Figure 6G-H). The varied additional positions observed for the Cys620 position in the other GAF2 homology models suggest there is flexibility in positioning of the cysteine residue. Specifically, the models based on 6OAQ and 6OAP place the cysteine residue so it is oriented away from the cofactor, and the model based on 6ZOH places it on the opposite side of the cofactor, near the C and D rings (Figure S12). To test the role of Cys620 in the Zn^2+^-induced fluorescence quenching, we mutated Cys to Ala in GAF2-PEB.

### Metal Sensitivity of GAF2_L652A_-PEB, GAF2_L652I_-PEB, and GAF2_C620A_-PEB Mutants

First, we examined the UV-visible absorption and fluorescence spectra of each mutant. GAF2_L652A_-PEB showed a decrease in absorbance and fluorescence (Φ = 0.14, Figure S4C-D, Table S2), which could be consistent with increased solvent access or flexibility of the bound bilin cofactor leading to a decrease in its brightness. The decrease in absorbance could also be a result of diminished binding efficiency of PEB during protein expression, so the fraction of bilin-bound protein to unbound protein is lower. The UV-visible absorption spectrum shows a peak at 492 nm, which is consistent with WT GAF2-PEB, but given the low absorbance any additional red-shifted peaks would likely be obscured. Attempts to measure the extinction coefficient were unsuccessful due to the lack of a clear signal and inability to unfold the protein without degrading the cofactor so that the concentration of PEB-bound protein relative to total protein could be estimated. The fluorescence emission maximum of GAF2_L652A_-PEB was blue-shifted to 553 nm from the 572 nm maximum observed for WT GAF2-PEB. GAF2_L652I_-PEB showed an increase in absorbance and decrease in fluorescence (Φ = 0.47, Figure S4E-F, Table S2) compared to WT GAF2-PEB (Figure 1B, Table S2). The increased absorbance could be due to the change in the bilin environment or increased binding efficiency of PEB. The UV-visible absorption spectrum of GAF2_C620A_-PEB shows a maximum at 544 nm, similar to WT GAF2-PEB (λ_max_ = 546 nm), but the blue-shifted shoulder around 495 nm is mostly diminished (Figure S4G, Table S2). Given that the peak at 495 nm has been attributed to GAF2-PUB and that at 546 nm to GAF2-PEB, this result suggests that little to no autoisomerization of PEB to PUB upon binding to GAF2_C620A_ has occurred (Figure 1A).^25^ The level of absorbance is similar for the mutant and WT bilin-bound proteins at similar protein concentrations. The fluorescence emission maximum of GAF2_C620A_-PEB is at 567 nm (Φ = 0.70, Figure S4H, Table S2), also similar to WT GAF2-PEB.

We tested the effects of several biologically relevant metals on the fluorescence of GAF2_L652A_-PEB and GAF2_L652I_-PEB. Zinc induced an ∼5-fold increase in the fluorescence of GAF2_L652A_-PEB at pH 7.1, similar to that observed for WT GAF2-PEB (Figure 7A). The fluorescence maximum of GAF2_L652A_-PEB was blue-shifted upon addition of Zn^2+^ from 553 to 517 nm (Figure 7D). A small apparent enhancement in the integrated fluorescence was observed for Cu^+/2+^ addition to GAF2_L652A_-PEB, which can be attributed to increased scatter detected (Figure S13A-B). As for GAF2-PEB, the initial fluorescence emission of GAF2_L652A_-PEB was quenched upon addition of copper. No other metals induced any significant effects on the fluorescence of GAF2_L652A_-PEB (Figure 7A). In the case of GAF2_L652I_-PEB, the Zn^2+^-induced fluorescence enhancement was also retained (Figure 7B), although to a lesser extent than for the wildtype protein, possibly supporting decreased access to the site due to sterics. Here, the fluorescence spectrum also showed a blue-shift in the maximum emission from 572 nm to 517 nm, but another prominent peak at 547 nm appeared along with a red-shifted band around 593 nm. The peak at 547 nm resembles that observed for Zn^2+^ addition to wildtype GAF2-PEB at pH 6.5 (Figure 5C). Both oxidation states of copper reduced the fluorescence of GAF2_L652I_-PEB (Figure S13C-D) and iron and cobalt also led to statistically significant reductions in fluorescence (Figure S14A-B). We also carried out a zinc titration with GAF2_L652I_-PEB to determine the binding affinity (Figure S15). Using ADA as the buffering chelator, we determined a single site dissociation constant of 1.5 µM, which is similar to, although it represents a slightly weaker affinity than observed for the WT GAF2-PEB (Table S3). In addition, we investigated the effect of pH on the fluorescence and zinc-induced fluorescence enhancement of GAF2_L652I_-PEB. As for WT GAF2-PEB, the initial integrated fluorescence decreased with increasing pH, but without a change in emission wavelengths (Figure S16A) and addition of zinc led to fluorescence enhancement at all pH values, ranging from ∼2-fold at pH 5.5 to ∼3-4-fold from pH 6.5-7.0 and up to ∼9-fold at pH 9.0 (Figure 8A). The level of reversibility also decreases with increasing pH, where the zinc-induced fluorescence is fully reversible through pH 7.5, and then partially reversible from pH 8.0-9.0. Compared to WT GAF2-PEB, GAF2_L652I_-PEB is reversible up to a higher pH value (7.5 versus 7.0). Examination of the fluorescence spectrum of this mutant in the presence of Zn^2+^ is consistent with the proposed protonation states discussed for WT GAF2-PEB. There is a shift in the wavelength of the shoulder present at ∼560 nm at pH 5.5 and 6.0 to 547 nm from pH 6.5 to 8.0 before it decreases (Figure 8B). In WT GAF2-PEB, this shoulder appears at pH 6.5 then decreases at pH 7.0 (Figure 5C). Given the autoisomerization of PEB to PUB, multiple bilin species are likely present in the binding site, but the shifting of spectra with pH to higher values for GAF2_L652I_-PEB compared to WT GAF2-PEB suggests that the p*K*_a_’s involved in zinc binding can be modulated via mutagenesis. Collectively, these results support the hypothesis that the Leu residue near the PEB binding site depicted in the GAF2 homology model can affect binding of Zn^2+^ to the bilin and impact the fluorescence properties.

**Figure 7.**
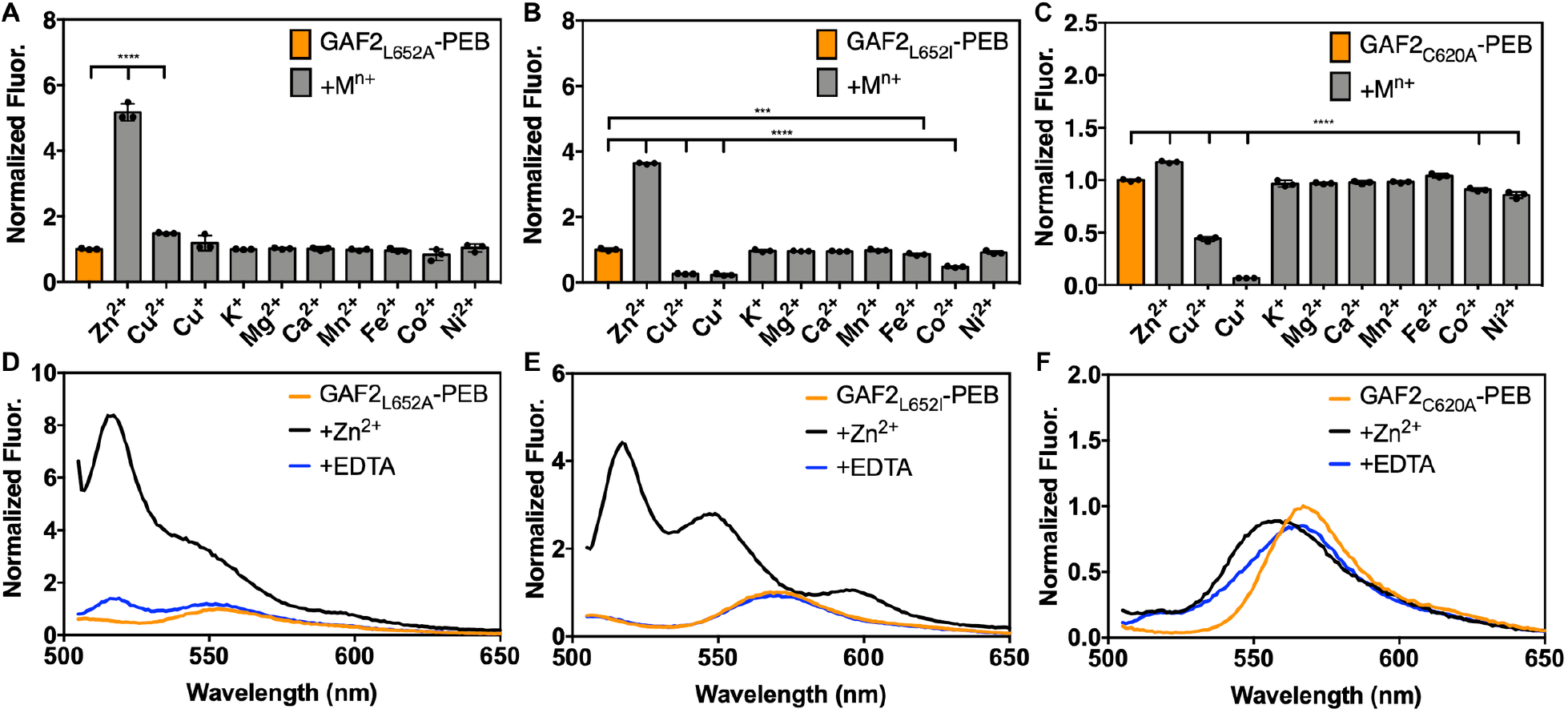
Metal sensitivity of mutant GAF2-PEB mutant proteins. Metal sensitivity of A) (His_6_)GAF2_L652A_-PEB, B) (His_6_)GAF2_L652I_-PEB, and C) GAF2_C620A_-PEB. To 3 μM of protein, 266 eq (800 µM K^+^, Mg^2+^, or Ca^2+^) or 33 eq (100 µM Zn^2+^, Cu^2+^, Cu^+^, Mn^2+^, Fe^2+^, Co^2+^, or Ni^2+^) metal salt was added. Data are shown as normalized integrated fluorescence emission (505-800 nm). λ_ex_ = 495 nm and λ_em_ = 505-800 nm. Data were analyzed by means of one-way analysis of variance (ANOVA); ***p < 0.001, ****p < 0.0001. Individual data points are overlaid on the bar chart representation. All error bars represent standard deviation for 3 replicates. D) Fluorescence emission spectra of (His_6_)GAF2_L652A_-PEB (3 µM) initially (orange line), upon addition of ZnCl_2_ (100 µM, black line), and with addition of EDTA (500 µM, blue line). E) Fluorescence emission spectra of (His_6_)GAF2_L652I_-PEB (3 µM) initially (orange), upon addition of ZnCl_2_ (100 µM, black), and with addition of EDTA (500 µM, blue). F) Fluorescence emission spectra of GAF2_C620A_-PEB (3 µM) initially (orange), upon addition of ZnCl_2_ (100 µM, black), and with addition of EDTA (500 µM, blue). Spectra were normalized to the initial fluorescence intensity of (His_6_)GAF2_L652A_-PEB at 553 nm (D), (His_6_)GAF2_L652I_-PEB at 572 nm (E), or GAF2_C620A_-PEB at 567 nm (F). Buffer: 50 mM HEPES, 100 mM NaCl, pH 7.1.

**Figure 8.**
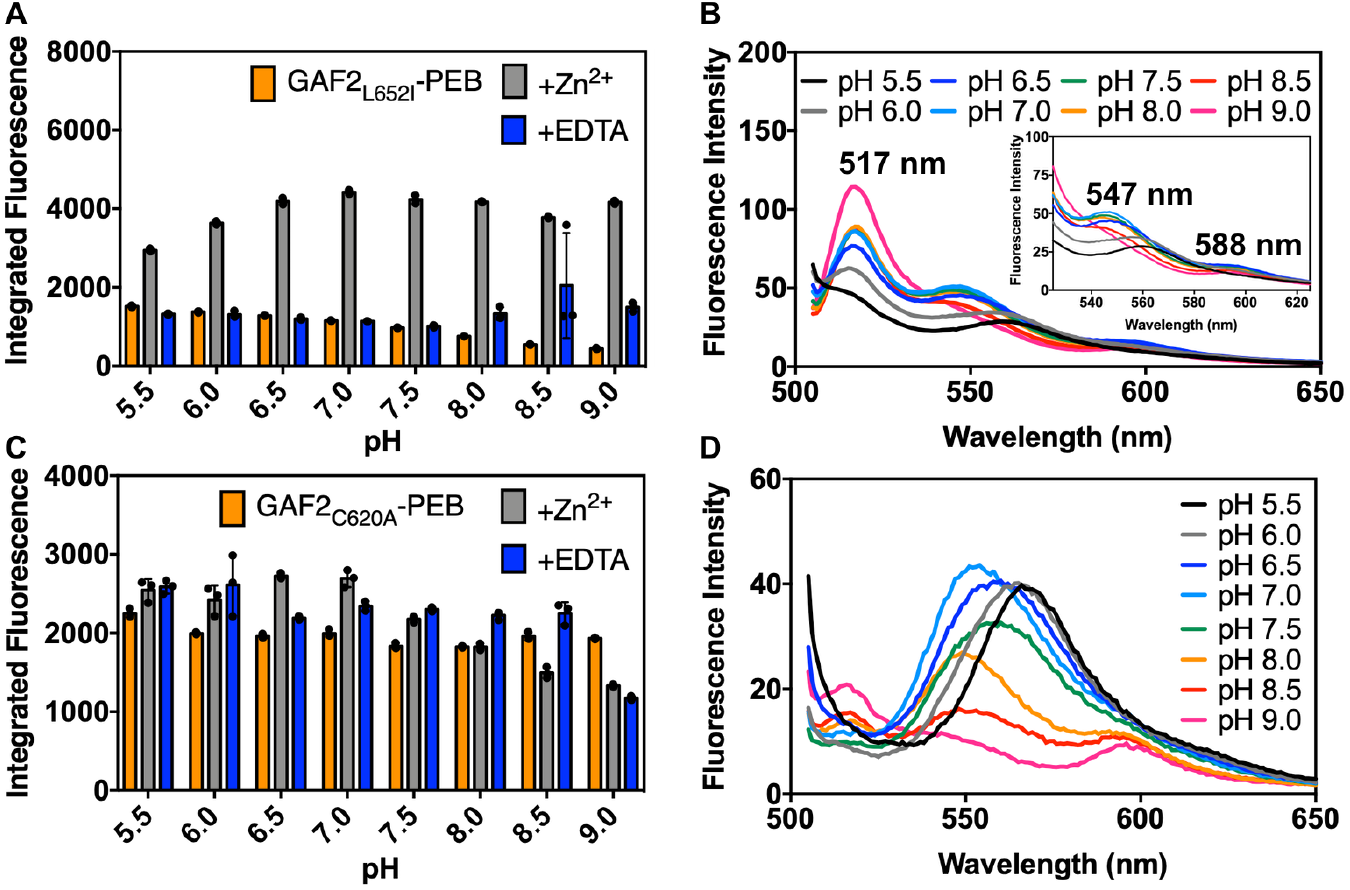
Effect of pH on the fluorescence intensity of A-B) GAF2_L652I_-PEB and C-D) GAF2_C620A_-PEB in the presence and absence of Zn^2+^. A, C) Integrated fluorescence intensity of GAF2-PEB mutant (orange, 3 μM), GAF2-PEB mutant with the addition of ZnCl_2_ (grey, 100 μM), and GAF2-PEB mutant + Zn^2+^ with the addition of EDTA (blue, 500 µM). λ_ex_ = 495 nm and λ_em_ = 505-800 nm. Individual data points are overlaid on the bar chart representation. All error bars represent standard deviation for 3 replicates. B) Spectra for GAF2_L652I_-PEB + Zn^2+^ representing the data shown in part A. Inset shows an expansion of the region from 525-625 nm. D) Spectra for GAF2_C620A_-PEB + Zn^2+^ representing the data shown in part C. Initial spectra and spectra with the addition of EDTA representing parts A and C are shown in Figure S16. Buffers: 50 mM MES, 100 mM NaCl (pH 5.5, 6.0, 6.5); 50 mM HEPES, 100 mM NaCl (pH 7.0, 7.5, 8.0); 50 mM CHES, 100 mM NaCl (pH 8.5, 9.0).

We also tested the effects of metal ions on the fluorescence of the DXCF motif cysteine mutant, GAF2_C620A_-PEB. Here, Zn^2+^ did not induce any substantial increase in the fluorescence intensity as was observed for WT GAF2-PEB and the leucine mutants (Figure 7C). The slight increase in fluorescence is even less than was observed for GAF3-PEB, which does not have a DXCF motif (Figure 2B). As for all of the other GAF proteins discussed thus far, Cu^2+/+^ both quench the fluorescence of GAF2_C620A_-PEB (Figure 7C, S13E-F). Compared to the other mutants and WT GAF2-PEB, Cu^2+^-induced fluorescence quenching is partially reversible rather than fully irreversible (Figure 2D, S13A,C,E). The Cu^+^-induced fluorescence quenching is more reversible, similar to GAF2_L652I_-PEB (Figure S13B,D,F). In addition, some small statistically significant decreases in fluorescence were observed with the addition of Co^2+^ and Ni^2+^, but there were no changes in the emission wavelengths (Figure 7C, S14C-D). To determine whether Zn^2+^ may still be binding to the bilin cofactor without leading to an increase in fluorescence as for WT GAF2-PEB, we examined the UV-visible absorption spectra. Here, no changes to absorption spectra were observed with addition of zinc and then EDTA, suggesting Cys620 is required for zinc coordination at the bilin cofactor (Figure S17). To determine whether the pH changes any zinc-induced fluorescence response in this mutant, we measured the fluorescence of GAF2_C620A_-PEB from pH 5.5-9.0 upon the addition of zinc and with subsequent addition of EDTA chelator (Figure 8C-D, S16C-D). Here, the initial fluorescence was not affected by pH (Figure S16C). This is distinct from WT GAF2-PEB where the initial fluorescence intensity generally decreased with increasing pH (Figure 5B). Addition of zinc leads to minor changes in the emission peak maximum and some decreases in fluorescence, but no overall increases in integrated fluorescence intensity (Figure 8C-D). Subsequent addition of EDTA led to the reversal of most spectra to the initial fluorescence spectra, with the exception of pH 9.0, which remained mostly quenched (Figure S16D). Collectively, these results suggest that the DXCF motif cysteine residue plays a crucial role in zinc-induced fluorescence enhancement in GAF2-PEB. It is possible that Cys620 may directly coordinate to bilin-bound Zn^2+^ within the protein, stabilizing a confirmation that leads to increased fluorescence. It is also possible that cysteine binding to the C10 position of the PEB and/or PUB cofactors facilitates Zn^2+^ binding to the cofactor (Figure 3B). A mixture of these possible binding configurations may be the source of the simultaneous blue- and red-shifts in fluorescence maxima for GAF2-PEB (Figure 2C).

## Conclusions

In summary, cyanobacteriochrome-derived fluorescent proteins are promising platforms for the design of new genetically encoded sensors that fluoresce across the visible to NIR spectrum, are relatively small, and offer the potential for studying oxygen-sensitive organisms. In this study, we report a novel turn-on zinc detection mechanism within a bilin-binding orange fluorescent protein and three single site mutants. The bilin-bound GAF2 domain from All1280 undergoes a reversible Zn^2+^-dependent fluorescence enhancement of ∼5-fold at pH 7, which increases with increasing pH. The 55 nm blue-shift in the fluorescence emission maximum suggests that there was a direct change in the chromophore, possibly due to direct metal binding. The simultaneous appearance of a small red-shifted shoulder suggests the presence of multiple species, possibly due to the partial autoisomerization of PEB to PUB within GAF2. Red-shifts in the UV-visible absorption maxima support an enhancement mechanism whereby Zn^2+^ can bind directly to the cofactor, likely at the pyrrole N atoms, as previously reported for free PEB.^43^ Titration with Zn^2+^ led to a binding curve that could be fitted to single site binding models with *K*_D_ = 1-5 µM at lower pH and a two-site binding model with *K*_D_ values of ∼20-80 nM at pH 8.0. Fluorescence from a similar PEB-binding GAF3 domain from Slr1393 was not enhanced with physiological concentrations of Zn^2+^ ions. Comparison of the crystal structure of GAF3 with a homology model of GAF2 reveals a possibly more solvent-accessible bilin cofactor for GAF2. Mutations around the solvent accessible pocket, including one that could sterically restrict access, led to changes in Zn^2+^-induced fluorescence enhancement in support of the hypothesis that Zn^2+^ can bind to PEB within the protein. More importantly, GAF2 harbors a DXCF motif with a cysteine residue near the bilin cofactor, which GAF3 does not have. Mutation of Cys620 to Ala confirms that this residue is crucial for Zn^2+^-induced fluorescence enhancement and the lack of any change to the UV-visible spectrum with zinc for this mutant confirms that this residue is also crucial for zinc binding. It is possible that Cys620 coordinates directly to zinc or that binding of Cys620 to the cofactor facilitates zinc binding. Collectively, this work provides insight to how bilin-binding fluorescent proteins may be harnessed to develop new metal ion sensors by utilizing direct metal binding to the bilin cofactor. Future work will aim to better understand metal interactions within the bilin binding site and to engineer higher affinity variants that could be used as zinc biosensors in living systems.

## Supporting Information

The Supporting Information includes cofactor structures, primers, additional characterizations of proteins, additional homology models, and the nucleotide and protein sequence of GAF2.

## Accession codes

Slr1393 GAF3: UniProt KB P73184, residues 441–596

All1280 GAF2: UniProt KB Q8YXD3, residues 562-727

GAF (*Spirulina subsalsa*): NBCI WP_017306776.1

PcyA: UniProtKB Q55891

HO1: UniProtKB P72849

PebS: UniProtKB Q58MU6

## Notes

The authors declare to no competing financial interest.

## Acknowledgement

We would like to acknowledge the funding support from NIH-NIGMS grant R35 GM138223 (M.L.Z.), The Welch Foundation grant E-1972 (M.L.Z.), and the University of Houston New Faculty Startup Grant (M.L.Z.).

