## Supporting Information for "Zinc-Induced Fluorescence Turn-on in Native and Mutant Phycoerythrobilin-Binding Orange Fluorescent Proteins"

### Table of Contents

Table S1. Primers sequences used in this work Spectroscopic properties of GAF-PEB proteins ........ S2

Table S2. Spectroscopic properties of GAF-PEB proteins ........ S2

Table S3. Zinc dissociation constants of GAF-PEB proteins ........ S2

Table S4. GAF2 homology model evaluation scores ........ S2

Scheme S1. Chemical structures of GAF-bound bilins in this work ........ S3

Figure S1. Nucleotide and amino acid sequence of the coding region of pET28a-GAF2 ........ S4

Figure S2. Example DynaFit fitting scripts for single site and two-site metal binding models ........ S5

Figure S3. SDS-PAGE of GAF-PEB proteins ........ S6

Figure S4. UV-visible and fluorescence spectra of GAF-PEB proteins ........ S7

Figure S5. Fluorescence spectra of GAF3-PEB proteins with zinc and copper ........ S8

Figure S6. Comparison of dose-dependent zinc sensitivity of $(His_6)$GAF2-PEB and $(His_6)$GAF3-PEB ........ S9

Figure S7. $Zn^{2+}$ binding titrations for GAF2-PEB, $(His_6)$GAF2-PEB, and $(His_6)$GAF3-PEB ........ S10

Figure S8. $Cu^{2+}$ binding titrations for GAF2-PEB, $(His_6)$GAF2-PEB, and $(His_6)$GAF3-PEB ........ S11

Figure S9. $Zn^{2+}$ binding titrations for GAF2-PEB at pH 6.5, 7.5, and 8.0 ........ S12

Figure S10. Comparison of GAF3 and GAF2 homology models ........ S13

Figure S11. Comparison of additional GAF2 homology models ........ S14

Figure S12. Comparison of Cys620 positioning across GAF2 homology models ........ S15

Figure S13. Fluorescence spectra of GAF2 mutant proteins with copper ........ S16

Figure S14. Fluorescence spectra of $(His_6)GAF2_{L652A}$-PEB with metals ........ S17

Figure S15. $Zn^{2+}$ binding titration for $(His_6)GAF2_{L652I}$-PEB ........ S18

Figure S16. Effect of pH on the fluorescence intensity of $GAF2_{L652I}$-PEB and $GAF2_{C620A}$-PEB ........ S19

Figure S17. UV-visible spectra of $GAF2_{C620A}$-PEB with zinc ........ S20

**Table S1. Primer Sequences Used in This Work.**

| Primers | Sequence |
|---|---|
| pET28a-GAF2_For | 5'- CCGACGGATCCATGCAATCATTGAATTTACAGAATATTCTGAACGC -3' |
| pET28a-GAF2_Rev | 5'- GCGACTTCTCGAGTTATGCCAGTTCGTTTTGC -3' |
| pET28a-GAF2$_{L652A}$_For | 5'- TCACCTGCGTGCGTTGGAGCAGTT -3' |
| pET28a-GAF2$_{L652A}$_Rev | 5'- CAGTCAGTCAGCTCAGCG -3' |
| pET28a-GAF2$_{L652I}$_For | 5'- TCACCTGCGTATTTTGGAGCAGTTC -3' |
| pET28a-GAF2$_{L652I}$_Rev | 5'- CAGTCAGTCAGCTCAGCG -3' |
| pET28a-GAF2$_{L652W}$_For | 5'- TCACCTGCGTTGGTTGGAGCAGT -3' |
| pET28a-GAF2$_{L652W}$_Rev | 5'- CAGTCAGTCAGCTCAGCG -3' |
| pET28a-GAF2$_{C620A}$_For | 5'- CCAGGATAATGCGTTCCAATCCGGCGCGGGC-3' |
| pET28a-GAF2$_{C620A}$_Rev | 5'- ATGTCTGCACCCAGCGCG-3' |

**Table S2. Spectroscopic Properties of GAF-PEB proteins.**

| Protein | Absorption | | Fluorescence | |
|---|---|---|---|---|
| | $\lambda_{max}$ (nm) | ε ($mM^{-1}$ $cm^{-1}$) | $\lambda_{max}$ (nm) | Φ |
| GAF2-PEB[a] | 495, 546 | 49, 24 | 572 | 0.68 |
| GAF2$_{L652A}$-PEB | 492 | NA[b] | 553 | 0.14 ± 0.01 |
| GAF2$_{L652I}$-PEB | 496, 545 | NA[b] | 572 | 0.47 ± 0.06 |
| GAF2$_{C620A}$-PEB | 544 | NA | 567 | 0.70 ± 0.03 |
| GAF3-PEB[a] | 530 | 43 | 575 | 0.31 |

a. Taken from reference [1]. b. The mutant extinction coefficients could not be accurately determined due to an inability to measure the concentration of PEB-bound protein relative to total protein because the protein could not be denatured without degrading the cofactor.

**Table S3. Zinc dissociation constants of GAF-PEB proteins.**

| Protein | pH | Buffering chelator | $K_D$ (M) |
|---|---|---|---|
| GAF2-PEB | 7.1 | ADA (1 mM) | 6.5 (0.1) × $10^{-7}$ |
| GAF2-PEB | 6.5 | ATP (1 mM) | 5 (1) × $10^{-6}$ |
| GAF2-PEB | 7.0 | ATP (1 mM) | 1.1 (0.1) × $10^{-6}$ |
| GAF2-PEB | 8.0 | ADA (1 mM) | 7.7 (0.8) × $10^{-8}$, 2.0 (0.7) × $10^{-8}$ |
| ($His_6$)GAF2-PEB | 7.1 | ADA (1 mM) | 5.2 (0.1) × $10^{-7}$ |
| ($His_6$)GAF3-PEB | 7.1 | ADA (1 mM) | 4.4 (0.4) × $10^{-8}$ |
| ($His_6$)GAF2$_{L652I}$-PEB | 7.1 | ADA (1 mM) | 1.5 (0.5) × $10^{-6}$ |

**Table S4. GAF2 homology model evaluation scores.**

| Template PDB ID | Sequence identity (%) | QMeanDisCo | GMQE |
|---|---|---|---|
| 6OAP | 49.67 | 0.75 ± 0.05 | 0.76 |
| 6OAQ | 49.67 | 0.75 ± 0.05 | 0.76 |
| 6MGH | 48.98 | 0.74 ± 0.07 | 0.70 |
| 7LSC | 46.26 | 0.73 ± 0.05 | 0.70 |
| 5ZOH | 41.03 | 0.71 ± 0.07 | 0.70 |
| 5DFX | 40.13 | 0.72 ± 0.05 | 0.69 |

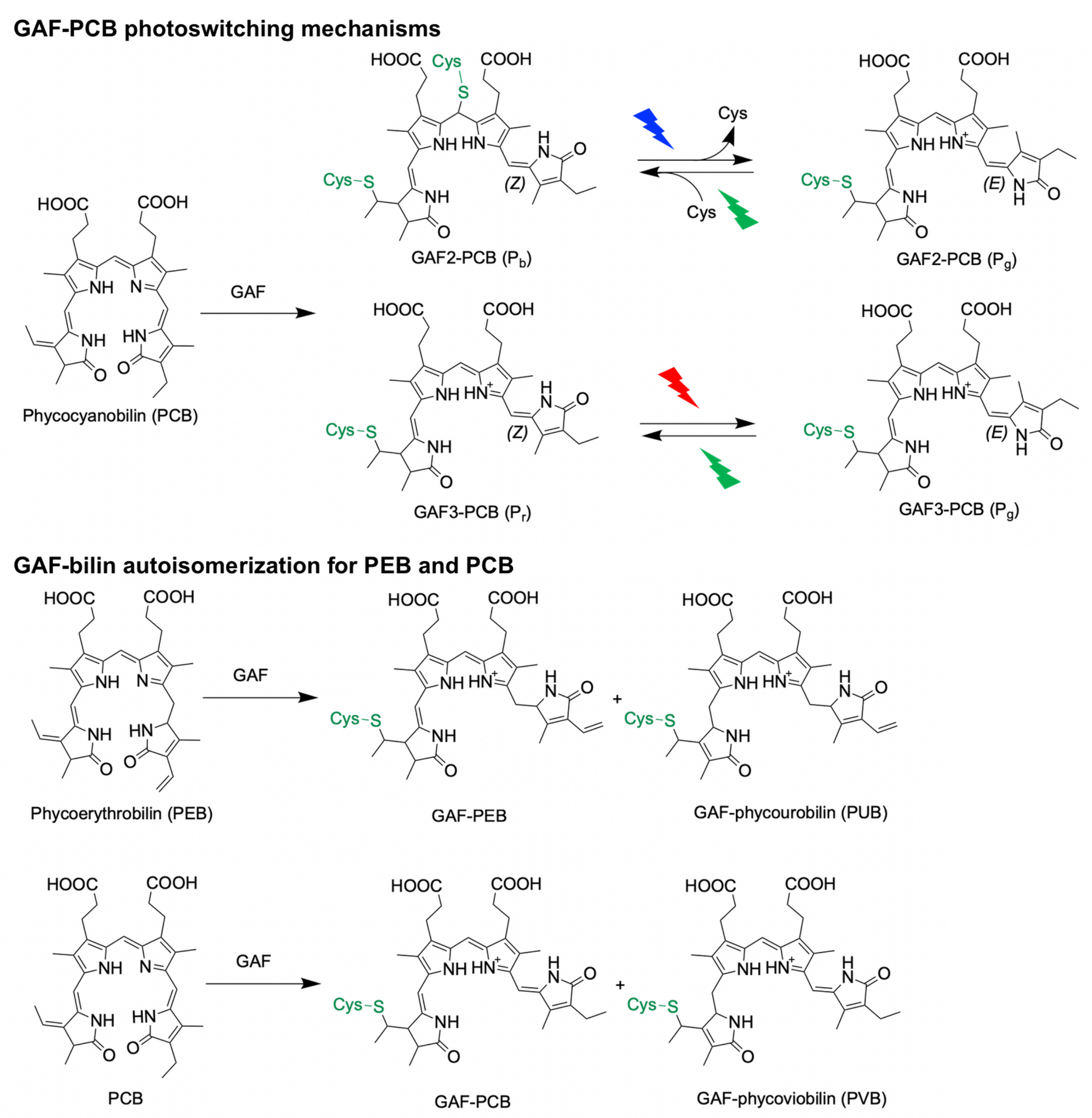


**Scheme S1. Chemical structures of GAF-bound bilins in this work.** Top: General photoswitching mechanisms for All1280 GAF2 (blue/green) and Slr1393 GAF3 (red/green) when bound to phycocyanobilin (PCB).[2–6] Bottom: Partial autoisomerization of GAF2/GAF3-phycoerythrobilin (PEB) to phycourobilin (PUB) occurs upon bilin binding.[1] Similarly, partial autoisomerization of GAF2-PCB to phycoviobilin (PVB) occurs upon bilin binding.[2,3]

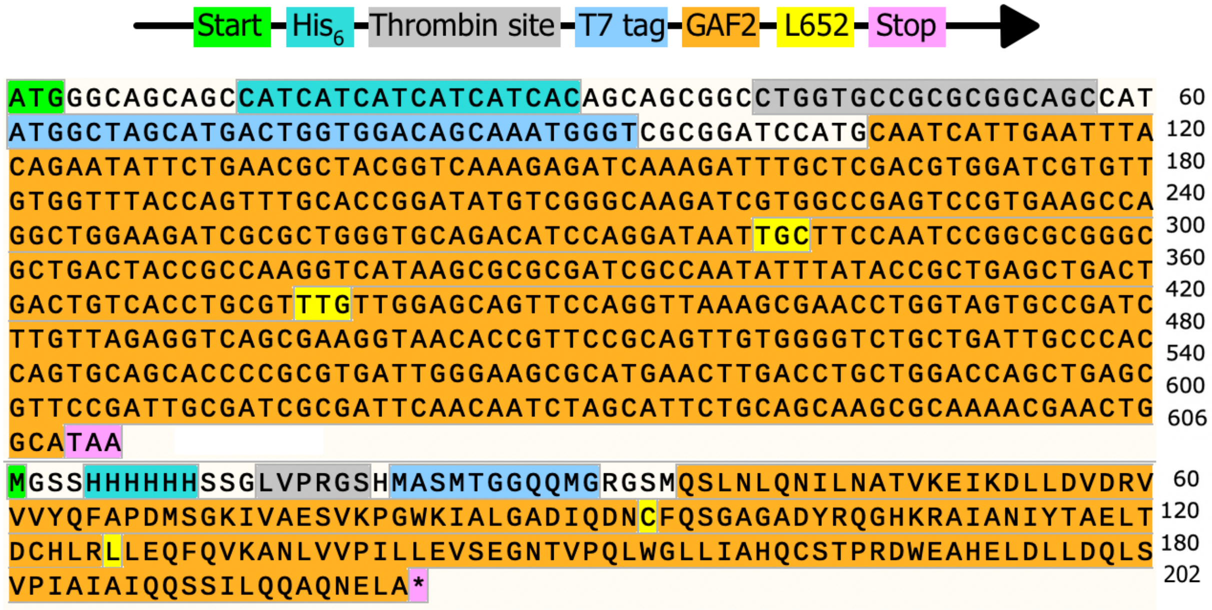


**Figure S1. Schematic (5' to 3') and nucleotide (top) and amino acid (bottom) sequences of the coding region of pET28a-GAF2.** The coding regions for the start codon, His6 tag, thrombin cleavage site, T7 tag, GAF2, and stop codon are all highlighted as shown. The coding sequence for the Cys residue at position 620 of All1280 (shown at position 94 in the amino acid sequence here) and for the Leu residue at position 652 of All1280 (shown at position 126 in the amino acid sequence here) of the GAF2 protein are colored yellow.

```
Fit binding affinity
[task]
   task = fit
   data = equilibria
[mechanism]
   s + m <==> s.m       :       kd1     dissociation
   e + m <==> e.m       :       kd2     dissociation
[constants]
   kd1 = 1e-10 ?
   kd2 = 9.4e-6
[concentrations]
   s = 3e-6
   e = 1e-3
[responses]
   s = 3e3 ?
   s.m = 4.6e7 ?
[data]
   variable m
   set data
[output]
   directory ./fittings/gafmutants/gaf2WT-Zn-ATP-pH-7-1

;_______________________________________________________

[set:data]

s        m
0       1
7.99E-05        3.299936656
1.60E-04        4.090389415
2.00E-04        4.788191764
2.39E-04        5.115938016
2.79E-04        5.249859411
3.19E-04        5.274524046
3.59E-04        5.219384138


[end]
```

```
Fit binding affinity
[task]
   task = fit
   data = equilibria
[mechanism]
   s + m <==> s.m       :       kd1     dissociation
   s.m + m <==> s.m.m   :       kd2     dissociation
   e + m <==> e.m       :       kd3     dissociation
[constants]
   kd1 = 1e-10 ?
   kd2 = 2e-8?
   kd3 = 6.44e-8
[concentrations]
   s = 3e-6
   e = 1e-3
[responses]
   s = 3e3 ?
   s.m = 4.6e7 ?
   s.m.m = 4.6e7 ?
[data]
   variable m
   set data
[output]
   directory ./fittings/gafmutants/gaf2WT-Zn-ADA-pH-8-1

;_______________________________________________________

[set:data]

s        m
0       1
7.99E-05        1.188395548
1.60E-04        1.412273565
2.00E-04        1.569223435
2.39E-04        1.789696311
2.79E-04        2.002303189
3.19E-04        2.235756177
3.59E-04        2.586970283
3.98E-04        2.90191875
4.38E-04        3.244136083
4.78E-04        3.55635032
5.17E-04        3.831096076
5.57E-04        4.133811401
5.96E-04        4.406673775
6.36E-04        4.621228037
6.75E-04        4.821201964
7.15E-04        5.030669494
7.54E-04        5.267881859
7.94E-04        5.499856246


[end]
```

**Figure S2.** Example DynaFit fitting scripts for a single site (left) and two-site (right) binding models. The model accounts for the chelator present to buffer metal ion concentrations. In the model "s" is the sensor, "m" is metal, and "e" is chelator (here it is ATP or ADA). In the single site script Kd1 is the dissociation constant being fitted and Kd2 is the known dissociation constant for the chelator. In the two-site script Kd1 and Kd2 are the dissociation constants being fitted and Kd3 is the known dissociation constant for the chelator.

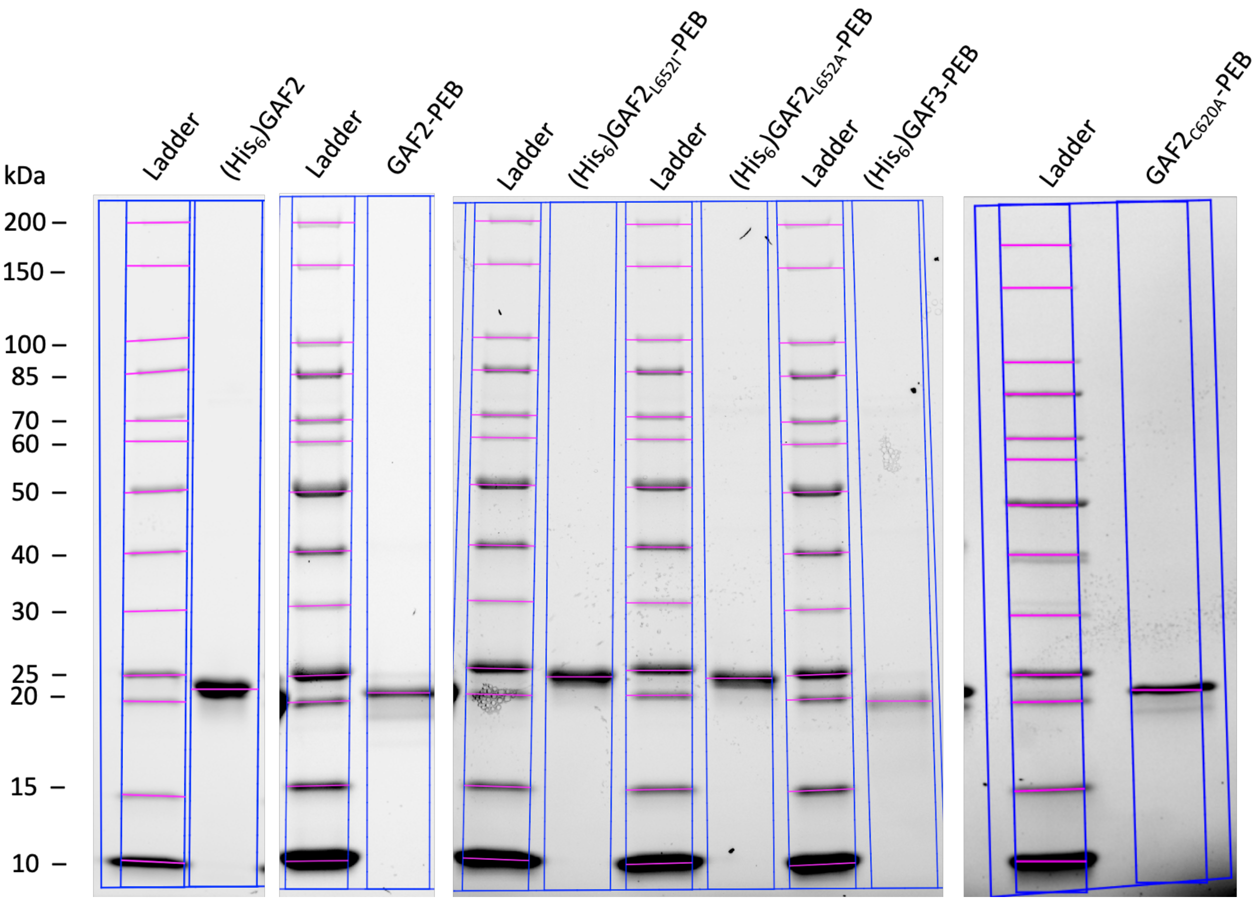


**Figure S3. SDS-PAGE of GAF-PEB proteins.** Denaturing polyacrylamide gel electrophoresis of purified GAF2-PEB, GAF3-PEB, and GAF2-PEB mutant proteins. Ladder: New England Biolabs Unstained Protein Standard, Broad Range, 10–200 kDa. Gel: BioRad 4–20% Mini-PROTEAN TGX Stain-Free Protein Gels.

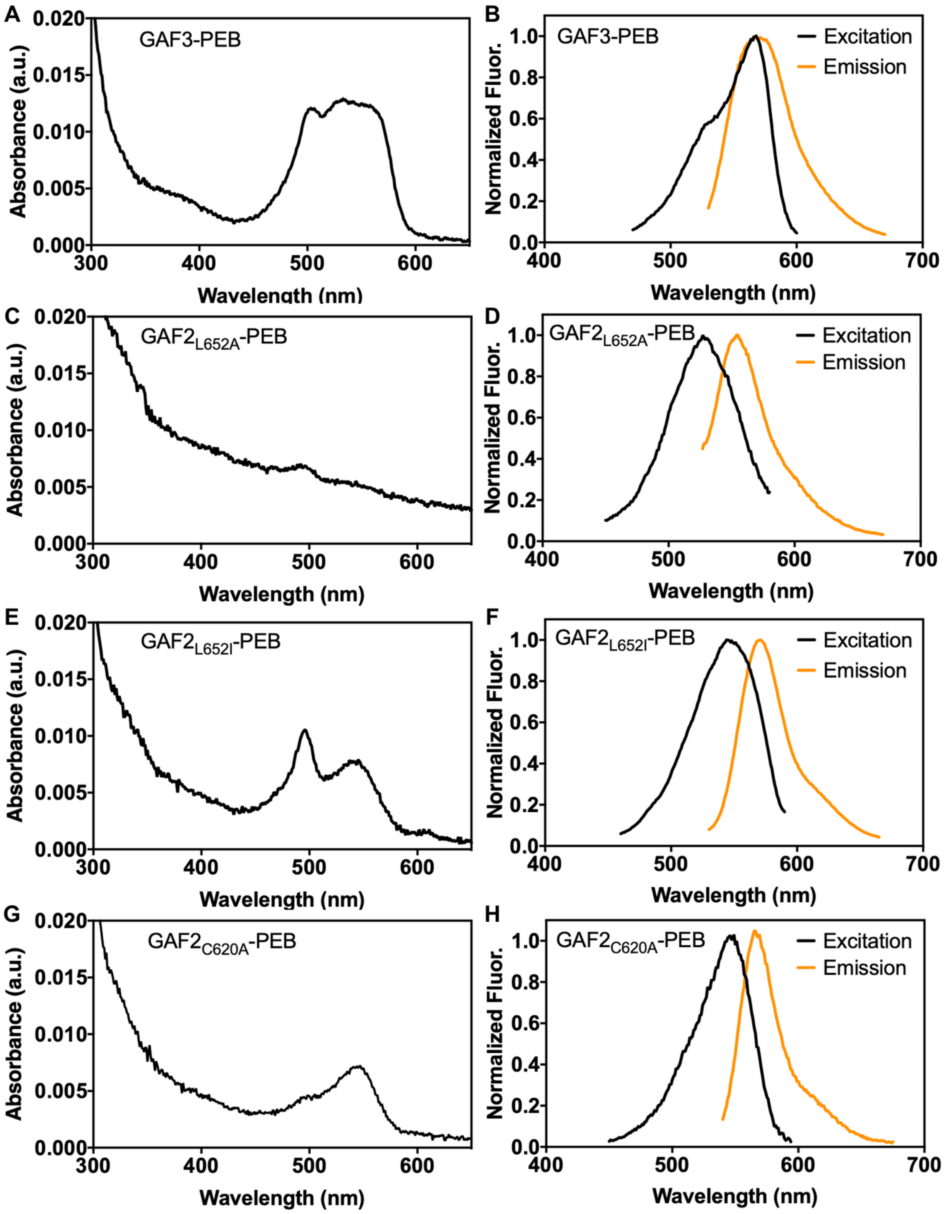


**Figure S4. UV-visible and fluorescence spectra of GAF-PEB proteins.** A) UV-visible spectrum of purified (His$_6$)GAF3-PEB (3.5 µM). B) Fluorescence excitation ($\lambda_{em}$ = 610 nm) and emission ($\lambda_{ex}$ = 520 nm) spectra of (His$_6$)GAF3-PEB. The excitation spectrum was normalized to the fluorescence at 568 nm. The emission spectrum was normalized to the fluorescence at 571 nm. C) UV-visible spectrum of purified (His$_6$)GAF2$_{L652A}$-PEB (5 µM). D) Fluorescence excitation ($\lambda_{em}$ = 600 nm) and emission ($\lambda_{ex}$ = 510 nm) spectra of (His$_6$)GAF2$_{L652A}$-PEB. The excitation spectrum was normalized to the fluorescence at 527 nm. The emission spectrum was normalized to the fluorescence at 553 nm. E) UV-visible spectrum of purified (His$_6$)GAF2$_{L652I}$-PEB (6.5 µM). F) Fluorescence excitation ($\lambda_{em}$ = 610 nm) and emission ($\lambda_{ex}$ = 510 nm) spectra of (His$_6$)GAF2$_{L652I}$-PEB. The excitation spectrum was normalized to the fluorescence at 545 nm. The emission spectrum was normalized to the fluorescence at 571 nm. G) UV-visible spectrum of purified GAF2$_{C620A}$-PEB (14 µM). H) Fluorescence excitation ($\lambda_{em}$ = 610 nm) and emission ($\lambda_{ex}$ = 495 nm) spectra of GAF2$_{C620A}$-PEB. The excitation spectrum was normalized to the fluorescence at 551 nm. The emission spectrum was normalized to the fluorescence at 571 nm. Buffer: 50 mM HEPES, 100 mM NaCl, pH 7.1.

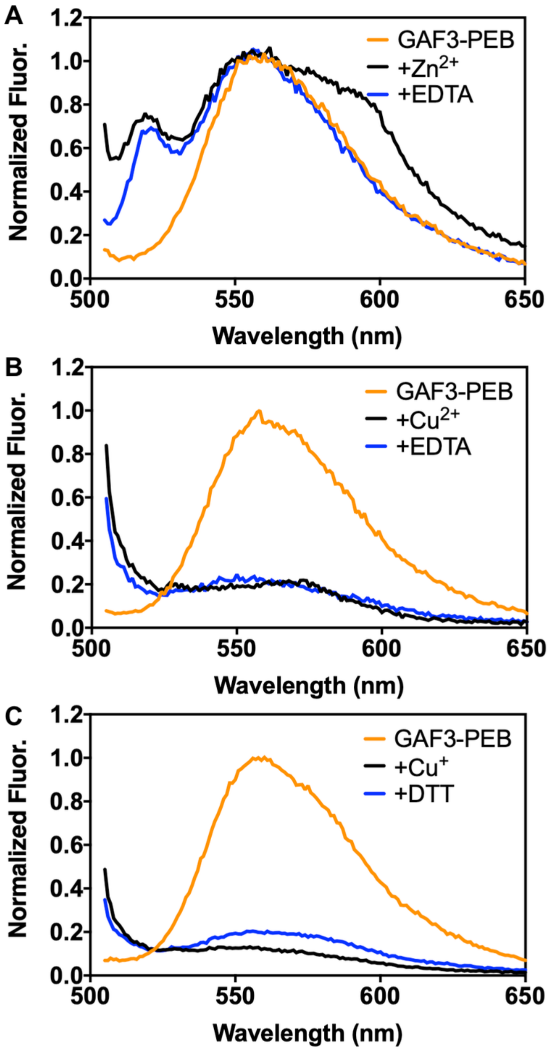


**Figure S5. Fluorescence spectra of (His$_6$)GAF3-PEB with zinc and copper.** A) Fluorescence emission spectra of (His$_6$)GAF3-PEB (3 µM) initially (orange line), upon addition of $ZnCl_2$ (100 µM, black line), and with addition of EDTA (500 µM, blue line). B) Fluorescence emission spectra of (His$_6$)GAF3-PEB (3 µM) initially (orange line), upon addition of $CuCl_2$ (100 µM, black line), and with addition of EDTA (500 µM, blue line). C) Fluorescence emission spectra of (His$_6$)GAF3-PEB (3 µM) initially (orange line), upon addition of $[Cu(CH_3CN)_4]PF_6$ (100 µM, black line), and with addition of DTT (200 µM, blue line). Fluorescence spectra were normalized to the maximum emission of GAF3-PEB at 557 nm. $\lambda_{ex}$ = 495 nm. Buffer: 50 mM HEPES, 100 mM NaCl, pH 7.1.

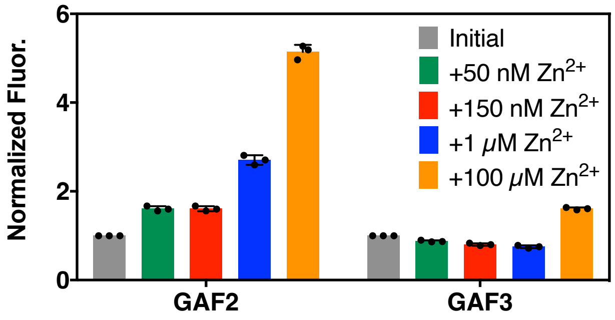


**Figure S6. Comparison of dose-dependent zinc sensitivity of (His$_6$)GAF2-PEB and (His$_6$)GAF3-PEB.** To 3 μM of protein, either 100 μM $ZnCl_2$ was directly added (excess, orange bar) or varied $ZnCl_2$ was added in the presence of 1 mM ADA (to buffer low concentrations of $Zn^{2+}$). $Zn^{2+}$ concentrations in the presence of 1 mM ATP (50 nM, 150 nM, and 1 μM) were calculated using MaxChelator. Data are shown as the integrated fluorescence normalized to the initial integrated fluorescence of the protein. $\lambda_{ex}$ = 495 nm and $\lambda_{em}$ = 505-800 nm. Individual data points are overlaid on the bar chart representation. Buffer: 50 mM HEPES, 100 mM NaCl, pH 7.1. All error bars represent the standard deviation for three replicates.

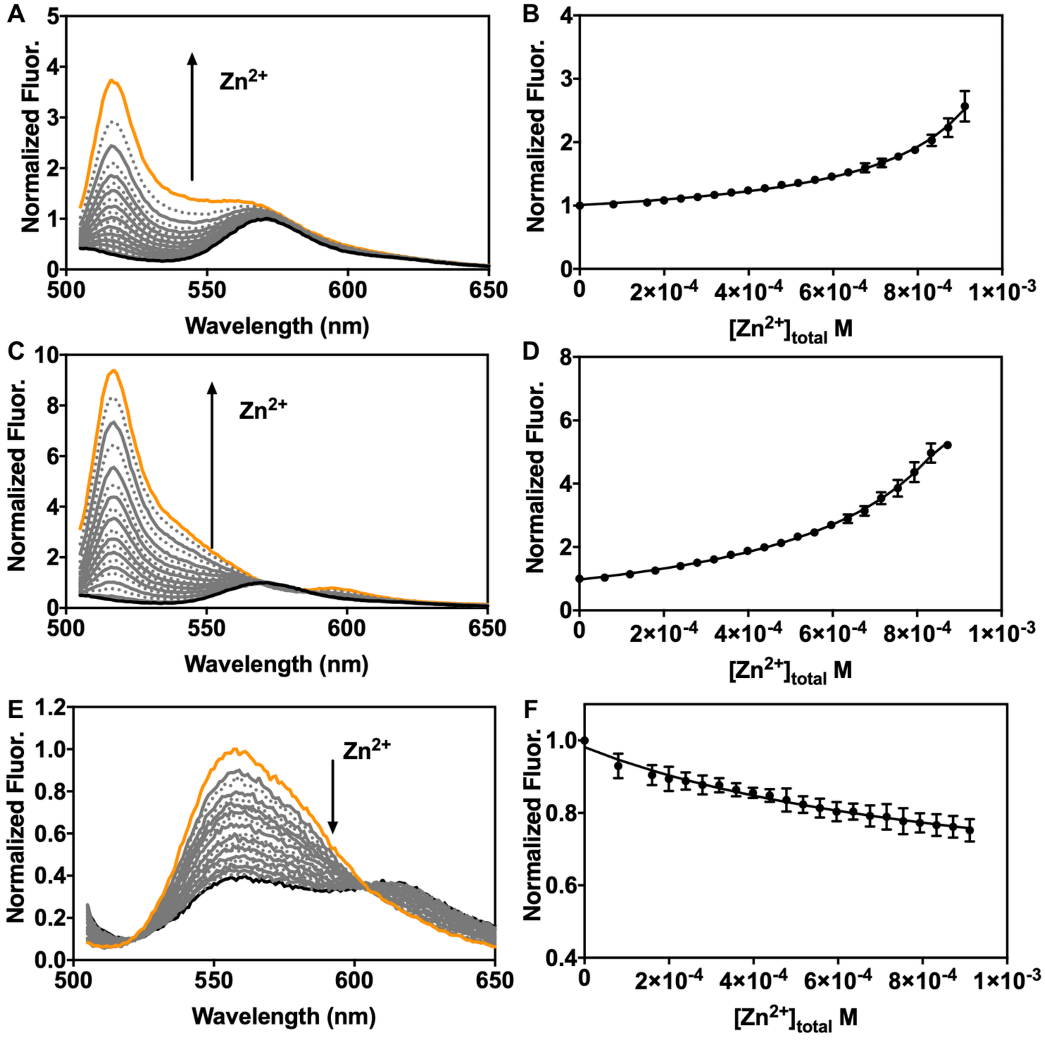


**Figure S7. $Zn^{2+}$ binding titrations for GAF2-PEB, (His$_6$)GAF2-PEB, and (His$_6$)GAF3-PEB in the presence of ADA.** A) Normalized fluorescence spectra of GAF2-PEB (3 μM) titrated with increasing amounts of $ZnCl_2$ in the presence of ADA (1 mM). B) Integrated fluorescence from part A plotted against total $Zn^{2+}$ concentration. Fluorescence was normalized relative to the initial GAF2-PEB fluorescence. C) Normalized fluorescence spectra of (His$_6$)GAF2-PEB (3 μM) titrated with increasing amounts of $ZnCl_2$ in the presence of ADA (1 mM). D) Integrated fluorescence from part C plotted against total $Zn^{2+}$ concentration. Fluorescence was normalized relative to the initial (His$_6$)GAF2-PEB fluorescence. E) Normalized fluorescence spectra of (His$_6$)GAF3-PEB (3 μM) titrated with increasing amounts of $ZnCl_2$ in the presence of ADA (1 mM). F) Integrated fluorescence from part E plotted against total $Zn^{2+}$ concentration. Fluorescence was normalized relative to the initial (His$_6$)GAF3-PEB fluorescence. All data were fitted with DynaFit (Methods Section). $\lambda_{ex}$ = 495 nm and $\lambda_{em}$ = 505-800 nm. Buffer: 50 mM HEPES, 100 mM NaCl, pH 7.1. All error bars represent the standard deviation for three replicates.

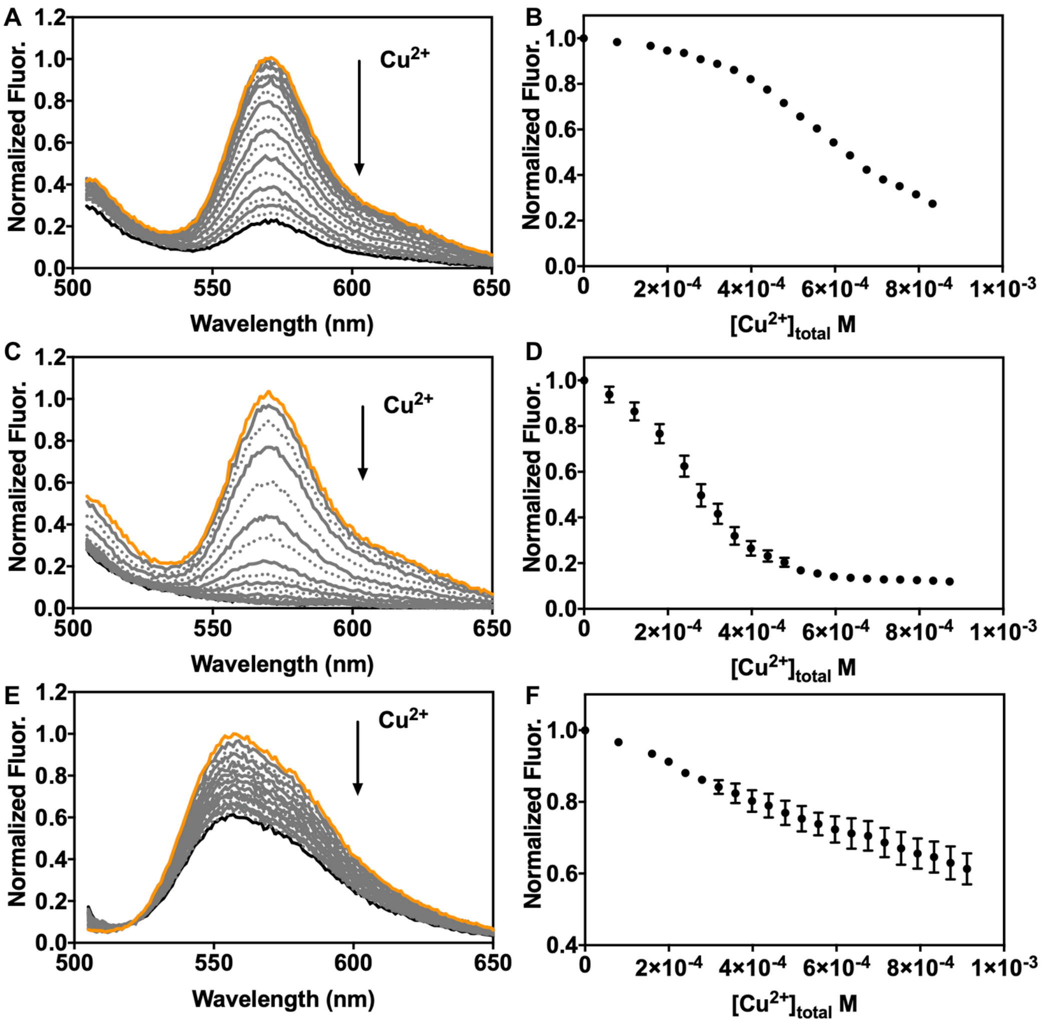


**Figure S8. $Cu^{2+}$ binding titrations for GAF2-PEB, (His$_6$)GAF2-PEB, and (His$_6$)GAF3-PEB in the presence of NTA.** A) Normalized fluorescence spectra of GAF2-PEB (3 μM) titrated with increasing amounts of $CuCl_2$ in the presence of NTA (1 mM). B) Integrated fluorescence from part A plotted against total $Cu^{2+}$ concentration. Fluorescence was normalized relative to the initial GAF2-PEB fluorescence. C) Normalized fluorescence spectra of (His$_6$)GAF2-PEB (3 μM) titrated with increasing amounts of $CuCl_2$ in the presence of NTA (1 mM). D) Integrated fluorescence from part C plotted against total $Cu^{2+}$ concentration. Fluorescence was normalized relative to the initial (His$_6$)GAF2-PEB fluorescence. E) Normalized fluorescence spectra of (His$_6$)GAF3-PEB (3 μM) titrated with increasing amounts of $CuCl_2$ in the presence of NTA (1 mM). F) Integrated fluorescence from part E plotted against total $Cu^{2+}$ concentration. Fluorescence was normalized relative to the initial (His$_6$)GAF3-PEB fluorescence. $\lambda_{ex}$ = 495 nm and $\lambda_{em}$ = 505-800 nm. Buffer: 50 mM HEPES, 100 mM NaCl, pH 7.1. All error bars represent the standard deviation for three replicates.

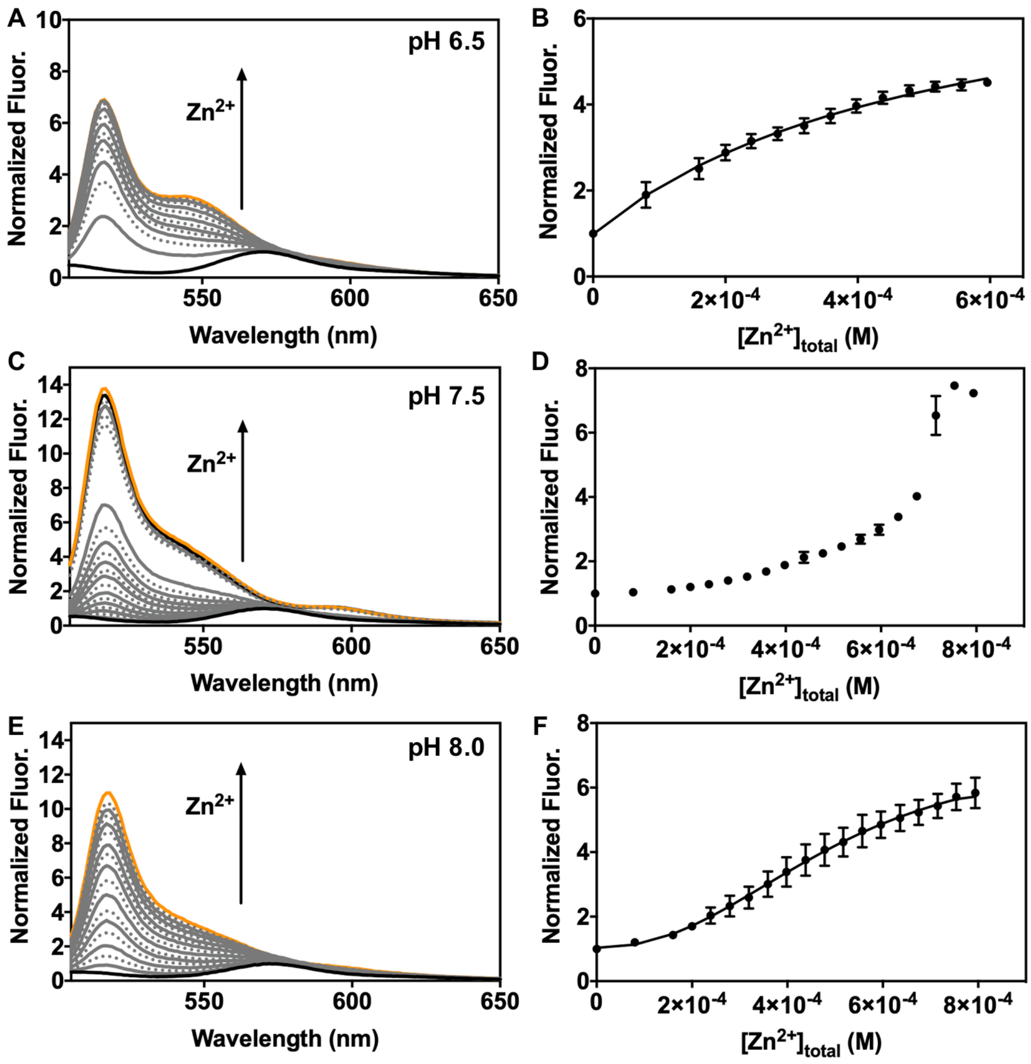


**Figure S9. $Zn^{2+}$ binding titrations for GAF2-PEB at varied pH.** A) Normalized fluorescence spectra of GAF2-PEB (3 μM) titrated with increasing amounts of $ZnCl_2$ in the presence of ATP (1 mM) at pH 6.5. B) Normalized integrated fluorescence from part A plotted against total $Zn^{2+}$ concentration. C) Normalized fluorescence spectra of GAF2-PEB (3 μM) titrated with increasing amounts of $ZnCl_2$ in the presence of ADA (1 mM) at pH 7.5. D) Normalized integrated fluorescence from part C plotted against total $Zn^{2+}$ concentration. E) Normalized fluorescence spectra of GAF2-PEB (3 μM) titrated with increasing amounts of $ZnCl_2$ in the presence of ADA (1 mM) at pH 8.0. F) Normalized integrated fluorescence from part E plotted against total $Zn^{2+}$ concentration. All fluorescence was normalized relative to the initial GAF2-PEB fluorescence at the given pH. Data were fitted with DynaFit (Materials and Methods Section and Figure SX). $\lambda_{ex}$ = 495 nm and $\lambda_{em}$ = 505-800 nm. Buffers: 50 mM MES, 100 mM NaCl (pH 6.5); 50 mM HEPES, 100 mM NaCl (pH 7.0, 7.5, 8.0). All error bars represent the standard deviation for three replicates.

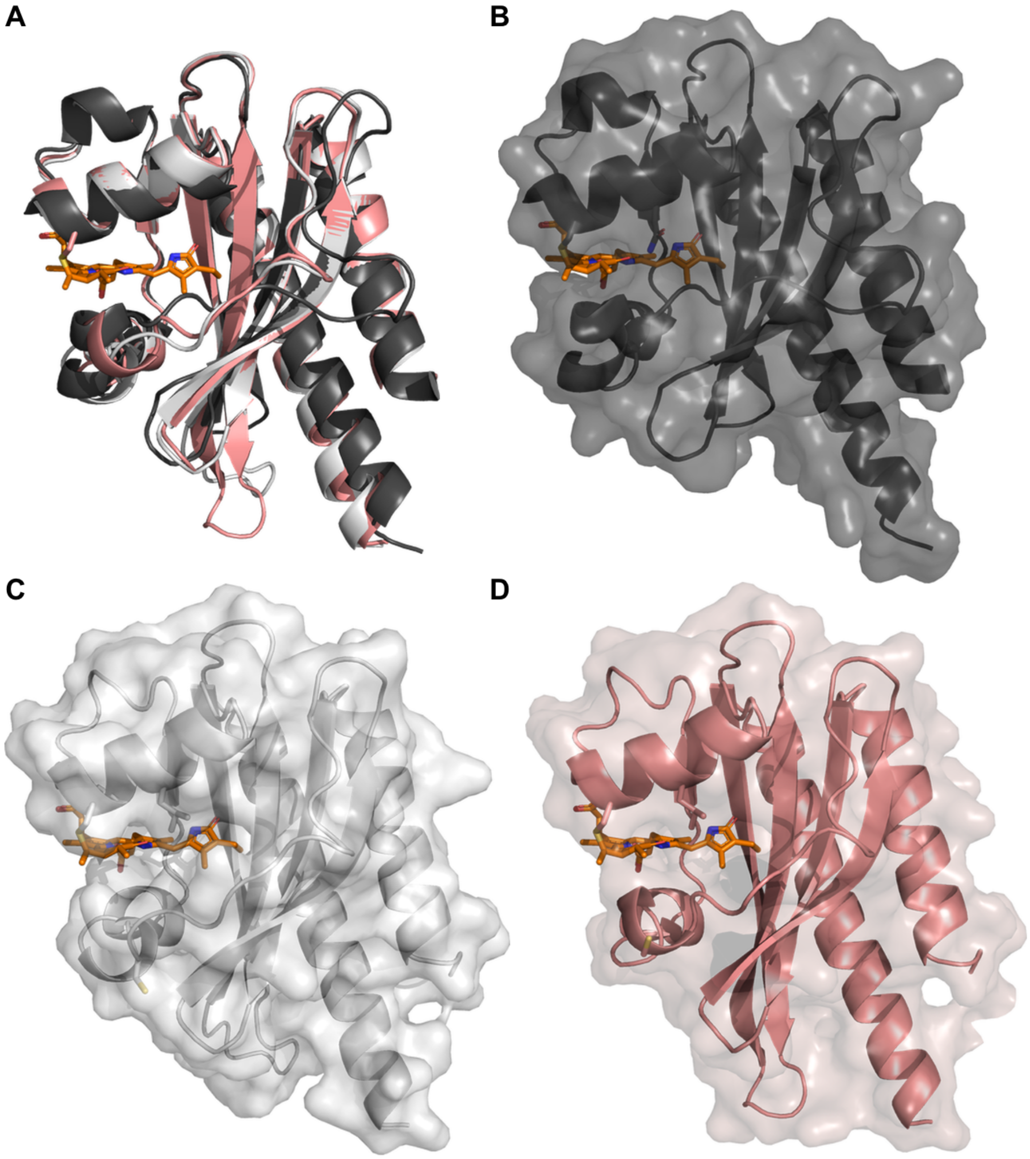


**Figure S10. Comparison of GAF3 and GAF2 homology models.** A) Overlay of the PyMOL representation of the crystal structure of Slr1393 GAF3 (black, PDB 5DFX), the homology model of All1280 GAF2 (grey, prepared using SwissModel and based on PDB 6OAP), and the homology model of All1280 GAF2 (pink, prepared using SwissModel and based on PDB 6OAQ). B) PyMOL model with surface representation for the 5DFX crystal structure of Slr1393 GAF3 (black, PDB 5DFX). C) PyMOL model with surface representation for the All1280 GAF2 homology model prepared using SwissModel and based on PDB 6OAP (grey). D) PyMOL model with surface representation for the All1280 GAF2 homology model prepared using SwissModel and based on PDB 6OAQ (pink). The bilin cofactor is colored orange and the Leu652 side chain is shown as a stick above the bilin cofactor in each model. The DXCF cysteine side chain is shown as a stick in a flexible loop region below the cofactor in part C-D.

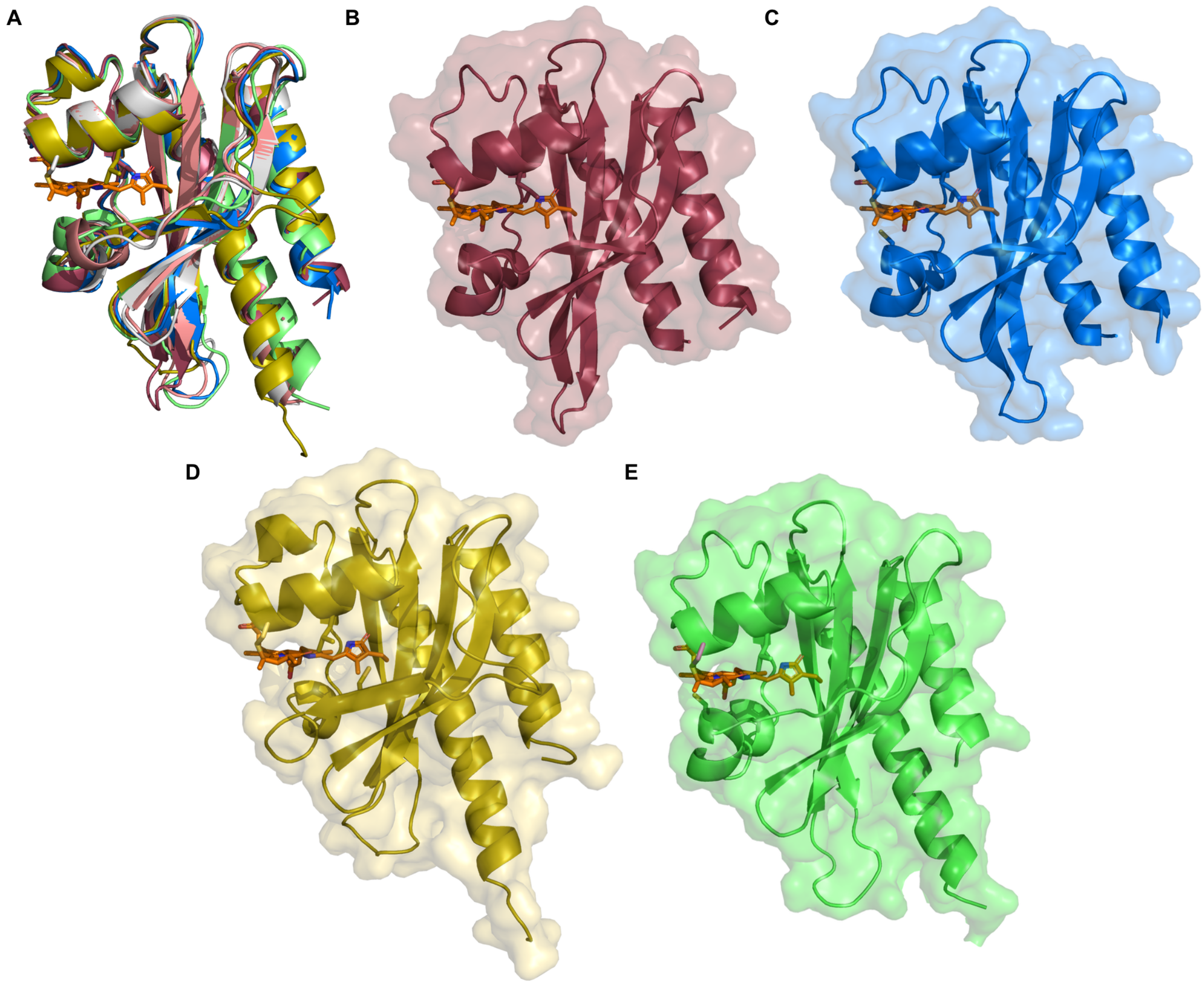


**Figure S11. Comparison of additional GAF2 homology models.** A) Overlay of the PyMOL representations of all All1280 GAF2 homology models described in this work. Homology models were prepared using SwissModel and are based on PDB 6OAP (grey), PDB 6OAQ (pink), PDB 6MGH (maroon), PDB 7LSC (blue), PDB 5ZOH (yellow), and PDB 5DFX (green). B) PyMOL model with surface representation for the All1280 GAF2 homology model prepared using SwissModel and based on PDB 6MGH (maroon). C) PyMOL model with surface representation for the All1280 GAF2 homology model prepared using SwissModel and based on PDB 7LSC (blue). D) PyMOL model with surface representation for the All1280 GAF2 homology model prepared using SwissModel and based on PDB 5ZOH (yellow). E) PyMOL model with surface representation for the All1280 GAF2 homology model prepared using SwissModel and based on PDB 5DFX (green). The bilin cofactor is colored orange and the Leu652 side chain is shown as a stick above the bilin cofactor in each.

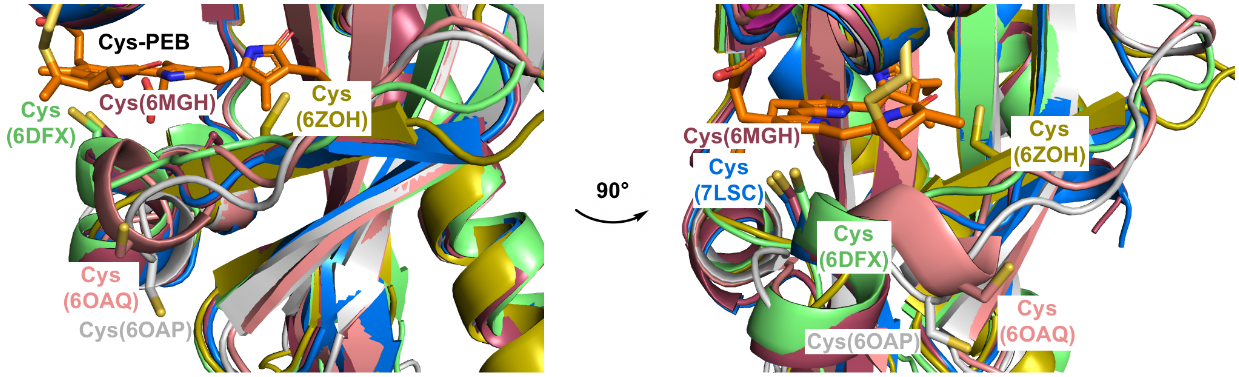


**Figure S12. Comparison of Cys620 positions across GAF2 homology models.** Two views of the overlay of the PyMOL representations for all All1280 GAF2 homology models. The Cys620 residue is shown as a stick in each homology model, which were prepared using SwissModel and are based on PDB 6OAP (gray), PDB 6OAQ (pink), PDB 6MGH (maroon), PDB 7LSC (blue), PDB 5ZOH (yellow), and PDB 5DFX (green). The bilin cofactor is colored orange labeled with Cys-PEB on the left view. The Cys620 residue is present in a flexible region of the protein and is found oriented in three general positions across the models. The models based on PDB 5DFX (green), PDB 6MGH (maroon), and PDB 7LSC (blue) all place the cysteine residue just below and closest to the cofactor. The models based on PDB 6OAQ (pink) and PDB 6OAP (gray) place the cysteine residue well below the cofactor and oriented away. Only the model based on PDB 6ZOH places the cysteine residue on a β sheet on the opposite side of the cofactor.

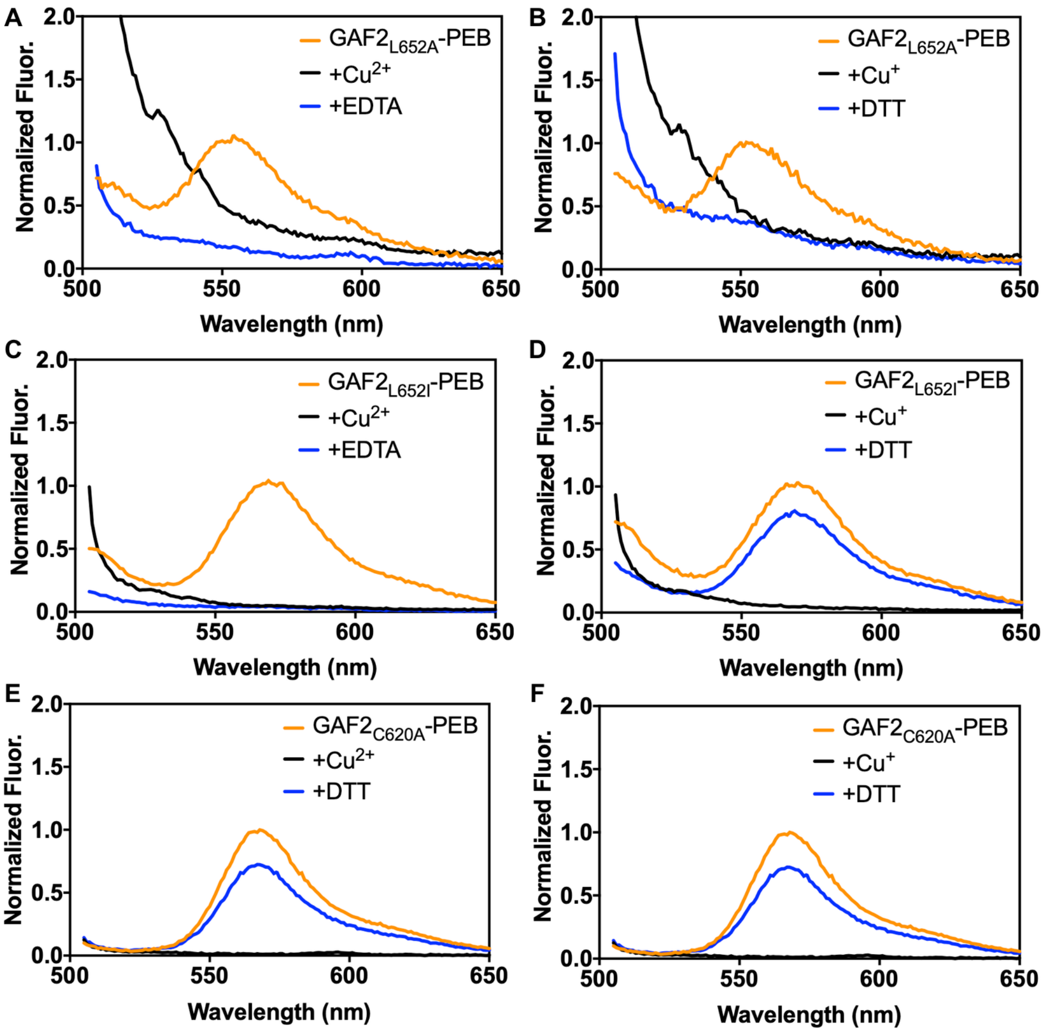


**Figure S13. Fluorescence spectra of GAF2 mutant proteins with copper.** A) Fluorescence emission spectra of $(His_6)GAF2_{L652A}$-PEB (4 μM) initially (orange line), upon addition of $CuCl_2$ (100 μM, black line), and with addition of EDTA (500 μM, blue line). B) Fluorescence emission spectra of $(His_6)GAF2_{L652A}$-PEB (4 μM) initially (orange line), upon addition of $Cu^+$ (100 μM, black line), and with addition of DTT (200 μM, blue line). C) Fluorescence emission spectra of $(His_6)GAF2_{L652I}$-PEB (4 μM) initially (orange line), upon addition of $CuCl_2$ (100 μM, black line), and with addition of EDTA (500 μM, blue line). D) Fluorescence emission spectra of $(His_6)GAF2_{L652I}$-PEB (4 μM) initially (orange line), upon addition of $Cu^+$ (100 μM, black line), and with addition of DTT (200 μM, blue line). E) Fluorescence emission spectra of $GAF2_{C620A}$-PEB (3 μM) initially (orange line), upon addition of $CuCl_2$ (100 μM, black line), and with addition of EDTA (500 μM, blue line). B) Fluorescence emission spectra of $GAF2_{C620A}$-PEB (3 μM) initially (orange line), upon addition of $Cu^+$ (100 μM, black line), and with addition of DTT (200 μM, blue line). Spectra were normalized to the initial fluorescence intensity of $(His_6)GAF2_{L652A}$-PEB at 553 nm (A-B), $(His_6)GAF2_{L652I}$-PEB at 572 nm (C-D), or $GAF2_{C620A}$-PEB at 567 nm (E-F). $\lambda_{ex}$ = 495 nm. Buffer: 50 mM HEPES, 100 mM NaCl, pH 7.1.

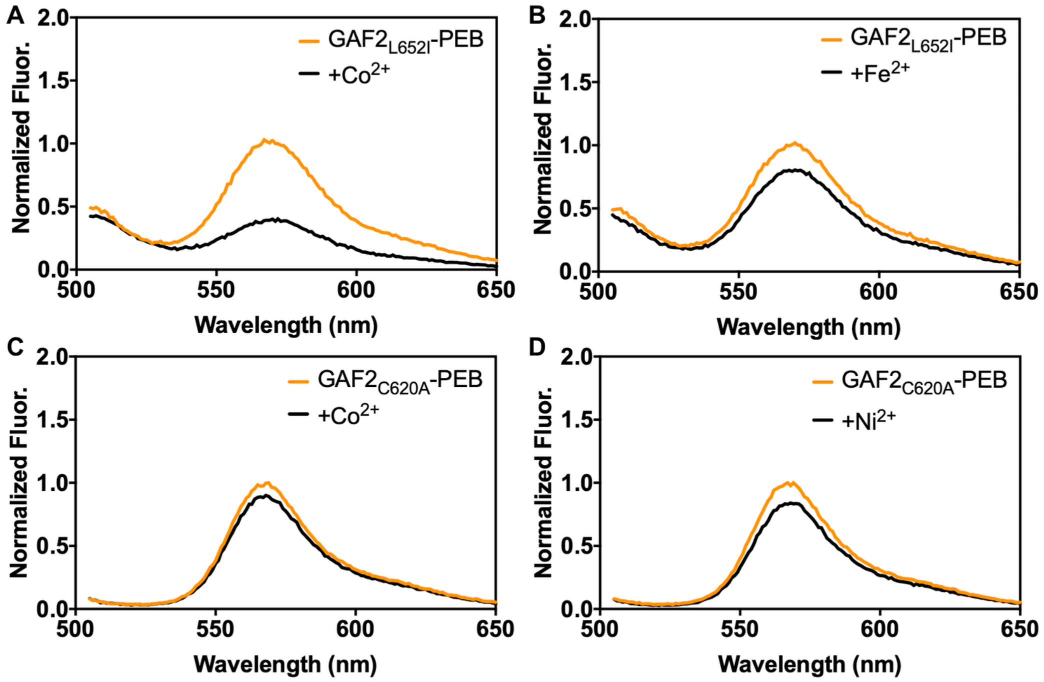


**Figure S14. Fluorescence spectra of (His$_6$)GAF2$_{L652I}$-PEB and GAF2$_{C620A}$-PEB with metals.** A) Fluorescence emission spectra of (His$_6$)GAF2$_{L652I}$-PEB (3 µM) initially (orange line) and upon addition of $CoCl_2$ (100 µM, black line). B) Fluorescence emission spectra of (His$_6$)GAF2$_{L652I}$-PEB (3 µM) initially (orange line) and upon addition of $FeCl_2$ (100 µM, black line). C) Fluorescence emission spectra of GAF2$_{C620A}$-PEB (3 µM) initially (orange line) and upon addition of $CoCl_2$ (100 µM, black line). D) Fluorescence emission spectra of GAF2$_{C620A}$-PEB (3 µM) initially (orange line) and upon addition of $NiCl_2$ (100 µM, black line). Fluorescence spectra were normalized to the maximum emission of (His$_6$)GAF2$_{L652I}$-PEB at 572 nm (A-B) or GAF2$_{C620A}$-PEB at 567 nm (C-D). $\lambda_{ex}$ = 495 nm. Buffer: 50 mM HEPES, 100 mM NaCl, pH 7.1.

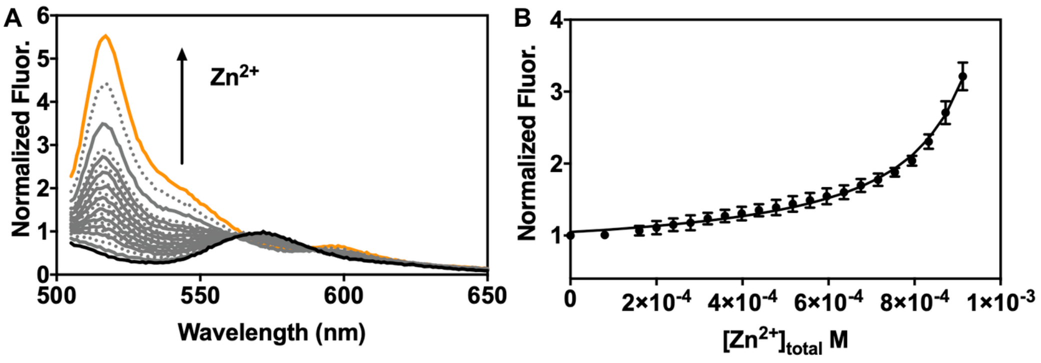


**Figure S15. $Zn^{2+}$ binding titration for (His$_6$)GAF2$_{L652I}$-PEB in the presence of ADA.** A) Normalized fluorescence spectra of (His$_6$)GAF2$_{L652I}$-PEB (3 μM) titrated with increasing amounts of $ZnCl_2$ in the presence of ADA (1 mM). B) Integrated fluorescence from part A plotted against total $Zn^{2+}$ concentration. Fluorescence was normalized relative to the initial (His$_6$)GAF2$_{L652I}$-PEB fluorescence. Data were fitted to a single site binding model with DynaFit (Methods Section). $\lambda_{ex}$ = 495 nm. $\lambda_{em}$ = 505-800 nm. Buffer: 50 mM HEPES, 100 mM NaCl, pH 7.1. All error bars represent the standard deviation for three replicates.

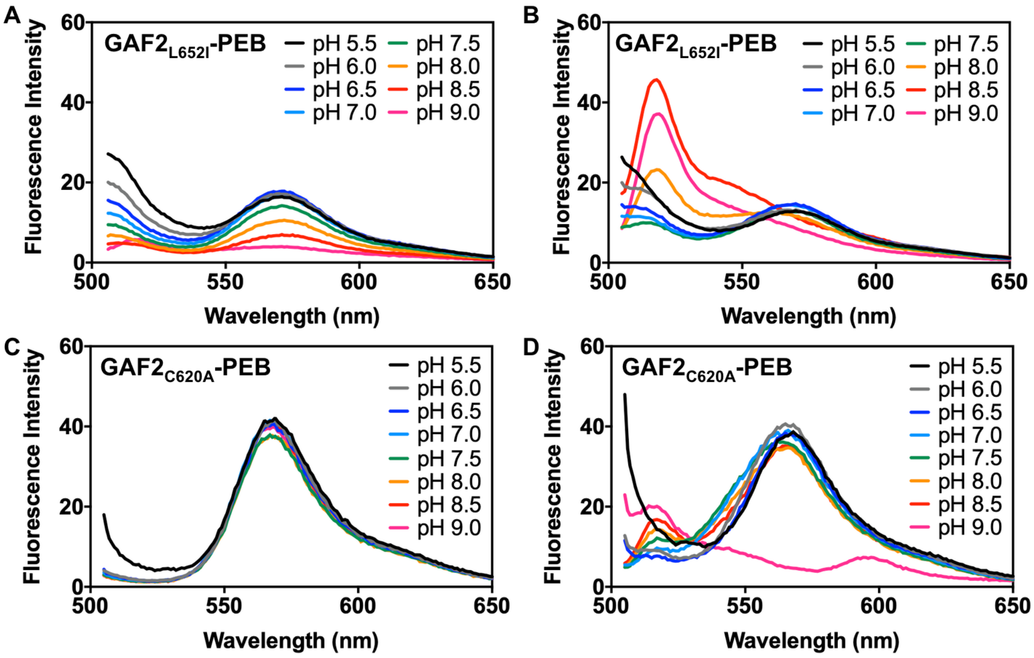


**Figure S16. Effect of pH on the fluorescence of A-B) GAF2$_{L652I}$-PEB and C-D) GAF2$_{C620A}$-PEB.** Spectra for A) GAF2$_{L652I}$-PEB (3 µM) and C) GAF2$_{C620A}$-PEB at varied pH. B) Spectra for GAF2$_{L652I}$-PEB + $ZnCl_2$ (100 µM) + EDTA (500 µM) at varied pH. D) Spectra for GAF2$_{C620A}$-PEB + $ZnCl_2$ (100 µM) + EDTA (500 µM) at varied pH. The spectra for each mutant and $ZnCl_2$ (100 µM) prior to the addition of EDTA are presented in Figure 8. The corresponding integrated fluorescence intensities for all spectra are also presented in Figure 8. Buffers: 50 mM MES, 100 mM NaCl (pH 5.5, 6.0, 6.5); 50 mM HEPES, 100 mM NaCl (pH 7.0, 7.5, 8.0); 50 mM CHES, 100 mM NaCl (pH 8.5, 9.0).

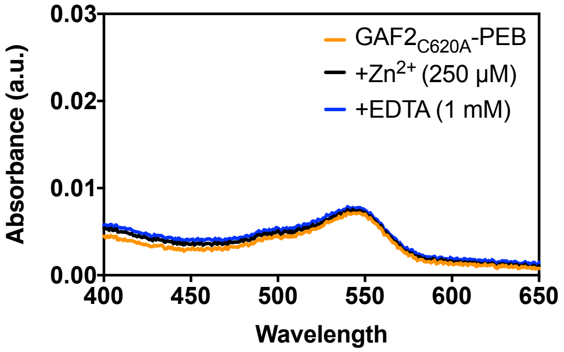


**Figure S17. UV-visible absorption spectra GAF2$_{C620A}$-PEB with zinc.** Absorption spectrum of GAF2$_{C620A}$-PEB (14 μM, orange line) initially, with addition of $ZnCl_2$ (250 μM, black line), and with addition of EDTA (1 mM, blue line). Buffer: 50 mM HEPES, 100 mM NaCl, 1 mM ATP, pH 7.1.
